# A multiomics approach to understanding pathology of Combined D,L-2- Hydroxyglutaric Aciduria and phenylbutyrate as potential treatment

**DOI:** 10.1101/2023.02.02.526527

**Authors:** Yu Leng Phua, Olivia M D’Annibale, Anuradha Karunanidhi, Al-Walid Mohsen, Brian Kirmse, Steven F Dobrowolski, Jerry Vockley

**Affiliations:** Department of Pediatrics, Division of Genetic and Genomic Medicine, UPMC Children’s Hospital of Pittsburgh, Pittsburgh, PA, USA; Department of Pathology, Clinical Biochemical Genetics Laboratory, UPMC Children’s Hospital of Pittsburgh, Pittsburgh, PA, USA; Department of Genetics and Genomic Sciences, Icahn School of Medicine at Mount Sinai, New York City, NY, USA; Department of Human Genetics, University of Pittsburgh Graduate School of Public Health, Pittsburgh, PA, USA; Department of Pediatrics, University of Mississippi Medical Center, Jackson, MS, USA

## Abstract

Combined D, L-2-Hydroxyglutaric Aciduria (D,L-2HGA) is a rare genetic disorder caused by recessive mutations in the *SLC25A1* gene that encodes the mitochondrial citrate carrier protein (CIC). *SLC25A1* deficiency leads to a secondary increase in mitochondrial 2-ketoglutarate that, in turn, is reduced to neurotoxic 2-hydroxyglutarate. Clinical symptoms of Combined D,L-2HGA include neonatal encephalopathy, respiratory insufficiency and often with death in infancy. No current therapies exist, although replenishing cytosolic stores by citrate supplementation to replenish cytosolic stores has been proposed. In this study, we demonstrated that patient derived fibroblasts exhibited impaired cellular bioenergetics that were worsened with citrate supplementation. We hypothesized treating patient cells with phenylbutyrate, an FDA approved pharmaceutical drug, would reduce mitochondrial 2-ketoglutarate, leading to improved cellular bioenergetics including oxygen consumption and fatty acid oxidation. Metabolomic and RNA-seq analyses demonstrated a significant decrease in intracellular 2-ketoglutarate, 2-hydroxyglutarate, and in levels of mRNA coding for citrate synthase and isocitrate dehydrogenase. Consistent with the known action of phenylbutyrate, detected levels of phenylacetylglutamine was consistent with the drug acting as 2-ketoglutarate sink in patient cells. Our pre-clinical studies suggest citrate supplementation is unlikely to be an effective treatment of the disorder. However, cellular bioenergetics suggests phenylbutyrate may have interventional utility for this rare disease.

## Introduction

Combined D,L-2- Hydroxyglutaric Aciduria (D,L-2HGA) is an ultra-rare genetic disorder caused by an autosomal recessive mutation in the *SLC25A1* gene that encodes the mitochondrial citrate carrier protein (CIC)^1,2^. Currently, ~20 individuals have been reported worldwide with this disorder, most with unique *SLC25A1* pathogenic mutations^1,3^. Symptoms of *SLC25A1* impairment include neonatal encephalopathy, respiratory insufficiency, developmental delay, hypotonia, and high mortality in infancy^1,3–6^. In spite of its rarity, Combined D,L-2HGA offers significant insights into the role of the intra-mitochondrial metabolic milieu in regulating energy metabolism. Functionally, the *SLC25A1* encoded mitochondrial citrate carrier (CIC) plays an indispensable role in maintaining mitochondrial citrate homeostasis, acting to export mitochondrial citrate into the cytosol in exchange for malate (Figure 1D)^1,3,4,7^. Impaired citrate export to the cytosol leads to excess in mitochondria with resultant secondary accumulation of 2-ketoglutarate that is further reduced into both D, and L-2-hydroxyglutarate, a recognized neurotoxin (Figure 1)^1^.

**Figure 1.**
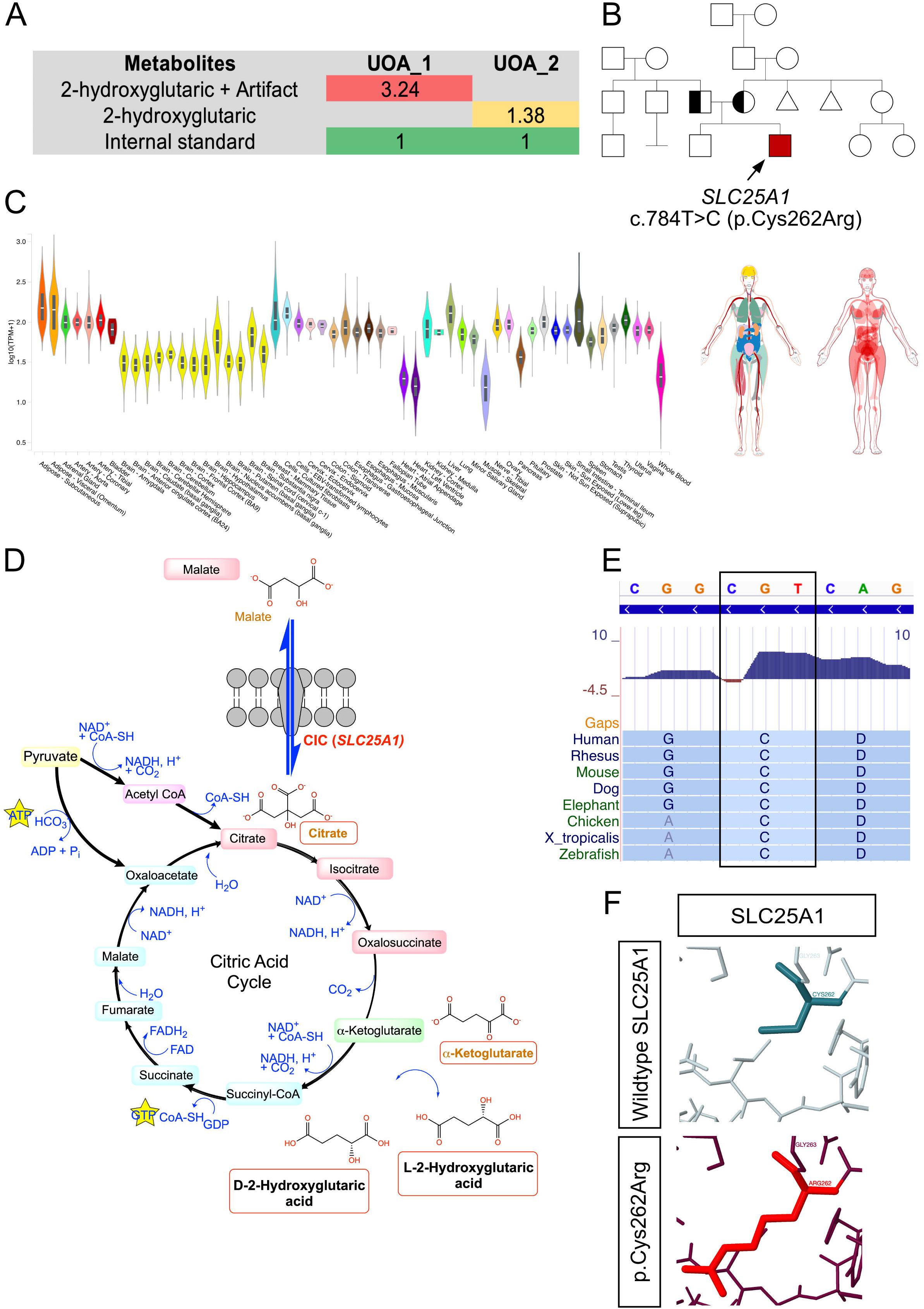
Biochemical and molecular analyses of a patient with Combined D,L-2HGA. **(A)** Gas chromatography mass spectrometry urine organics analysis of patient 893 urine specimen on 2 separate assessments revealed excessive secretion of 2-hydroxyglutarate. The metabolite values are qualitative ratios and are normalized to the internal spiked standard. **(B)** Whole exome sequencing identified a homozygous missense c.784T>C (p.Cys262Arg) variant in the *SLC25A1* gene in the proband, and Sanger sequencing verified both parents to be carriers of the variant. **(C)** GTEx Portal and The Human Protein Atlas databases revealed that *SLC25A1* is ubiquitously expressed in human tissues. **(D)** *SLC25A1* encodes for the citrate carrier protein (CIC) that acts to regulate the overall homeostasis of mitochondrial citrate through the exchange with cytoplasmic malate. The variant results in a loss of CIC channel activity which causes intramitochondrial accumulation of citrate that leads to secondary accumulation of 2-ketoglutarate that subsequently reduces into D- and L-2-Hydroxyglutaric acid which is neurotoxic. **(E)** Conservation analysis of the normal SLC25A1 protein cysteine residue at position 262 revealed shows it to be evolutionary conserved. **(F)** Structural modeling of the p.Cys262Arg variant predicts lack of structural impact on the integrity of the CIC channel.

Aside from the accumulation of metabolites causing various forms of toxicity, the pathomechanism of these toxicities is unclear, hampering efforts to develop effective therapies. D,L-2- Hydroxyglutarate has been postulated to impair mitochondrial energy metabolism, suggesting the need to reduce this nonphysiologic metabolite. The resulting decrease of citrate in the cytoplasm could alter complex lipid synthesis, including neural lipids, leading to the devastating neurodevelopmental impairment seen in patients. To address the latter possibility, therapeutic citrate supplementation has been suggested as a potential therapy. There is one report of treating genetically confirmed Combined D,L-2HGA patient with malate and citrate^4^. While oral administration of malate at 400 mg/kg/day failed to improve clinical presentation, oral citrate at 200-1500 mg/kg/day reduced seizures and D,L-2- Hydroxyglutarate excretion, but urinary 2-ketoglutarate excretion was concomitantly increased^4^. In a second patient, administration of 3mg/kg/day of oral citrate led to reduced frequency of apneic attacks, increased spontaneous respiratory drive, and transient blood lactate reduction^6^. However, D,L-2- Hydroxyglutarate remained markedly increased and the patient’s neurologic status remained poor^6^. The relatively modest response to citrate therapy was also mirrored in *in vitro* studies, whereby citrate supplementation in the patient fibroblasts resulted in reduced spare respiratory capacity as measured by micro-oximetry^6^.

In this study, we present *in vitro* fibroblast studies and clinical observations that citrate therapy for Combined D,L-2HGA is not helpful, but rather was detrimental in one patient. Alternatively, we hypothesized that sodium phenylbutyrate, an FDA-approved medication for the treatment of urea cycle deficiencies, maybe beneficial by reduction of intramitochondrial 2-ketoglutarate^8,9^. Metabolomic analysis of patient derived cells treated with phenylbutyrate points to significant reduction of 2-ketoglutarate and 2-hydroxyglutate levels through formation of phenylacetylglutamine (PAG), concomitant with improvement in cellular bioenergetics. Treatment of two Combined D,L-2HGA patients with phenylbutyrate confirmed accumulation of the non-toxic PAG in blood of both patients, and one patient presented improvement in clinical outcomes. These observations support the utility of repurposing of phenylbutyrate for treatment of Combined D,L-2HGA.

## Materials and Methods

The detailed materials and methods are available in the supplementary data

### Patient fibroblasts

Dermal punch biopsies enabled cell line establishment from molecular confirmed D,L-2HGA patients that were diagnosed through whole exome sequencing conducted in an approved CLIA-certified clinical lab. These fibroblast cell lines include, FB893 [NM_005984.5(*SLC25A7*):c.784T>C (p.Cys262Arg); Homozygous] and FB897 [NM_005984.5(*SLC25A7*):c.844C>T (p.Arg282Cys); c.821+1G>A] Variant phasing was confirmed through parental genotyping. Control fibroblasts from healthy individuals were functionally validated and reported in our previous study^10^. All experimental human protocols were approved by the Institutional Review Board at the University of Pittsburgh, protocol #19090211. Brief patient descriptions are presented in the Results.

### Cell culture and treatments

Fibroblasts were cultured and maintained in T175 flasks (Greiner Bio-One #660160) in complete Dulbecco’s Modified Eagle Medium media (Corning Life Sciences #10-013-CM) supplemented with 10% fetal bovine serum (ThermoFisher Scientific #26140079), 4 mM glutamine (ThermoFisher Scientific #25030081), 100 μg/mL Normocin anti-mycoplasma (InvivoGen #ant-nr-1), 100 IU penicillin and 100 μg/mL streptomycin (ThermoFisher Scientific #15140122) in a humidified 37 °C, 5% CO_2_ incubator. Routine mycoplasma screening was performed with the use of a Mycoplasma PCR Detection Kit (abm #G238). Sodium citrate (Millipore Sigma #71635) was reconstituted in PBS and used at a final concentration of 150 μM in complete DMEM media. 4-Phenylbutyric acid (Millipore Sigma #P21005) was first reconstituted in sterile MilliQ water, adjusted to pH 7.2 and used at a final concentration of 8 mM in complete DMEM media. The passage number of the fibroblasts used in the experiments were between 3-5 and treatments (citrate, phenylbutyrate) were performed for 72 hrs.

## Results

### Identification of a patient with Combined D,L-2- Hydroxyglutaric Aciduria

A 3-month-old male infant (Patient 893) first presented at UPMC Children’s Hospital of Pittsburgh (CHP) with a chief concern for hypotonia, apnea and feeding issues starting at 7 weeks of age. Progressively, the patient developed developmental delay, seizures, profound swallowing dysfunction and aspiration. Urine organics acid assessment at the CHP Clinical Biochemical Genetics Lab identified overrepresentation of 2-hydroxyglutaric acid. The urine 2-hydroxygluraic acid persisted in follow-up samples, and clinical whole exome testing performed at Baylor Genetics, Houston, TX, identified a novel homozygous NM_005984.5(*SLC25A1*):c.784T>C (p.Cys262Arg) variant of unknown significance (VUS) (Figure 1B). Mutations in *SLC25A1* are casual for Combined D,L-2- Hydroxyglutaric Aciduria (D,L-2HGA), an ultra-rare autosomal recessive neurometabolic disorder^1,3,4,6^. Sanger sequencing of the patient’s parents verified their carrier status. Given the VUS status of the c.784T>C (p.Cys262Arg) variant, we performed an *in vitro* citrate flux functional assay on FB893 fibroblasts which confirmed a >50% loss in CIC activity. These results have previously been reported^3^.

A second female patient (Patient 897) was identified with generalized seizures at 3 days of age at the University of Mississippi Medical Center. Initial EEG demonstrated global slowing and discontinuity suggesting cerebral dysfunction with reduced seizure threshold, but no active seizures. Brain MRI showed moderate to severe supratentorial ventriculomegaly with irregular margins of the lateral ventricles. The corpus callosum was markedly thinned and there was a large defect in the septum pellucidum (Supplemental Figure 1). Bilateral hippocampal incomplete inversion was noted. Metabolic evaluation identified the presence of 2-hydroxyglutarate in urine. Molecular diagnostic testing in a CLIA certified laboratory revealed two *SLC25A1* mutations in *trans*: c.844C>T (p.Arg282Cys) (Pathogenic) and c.821+1G>A (Splicing; Likely pathogenic). The patient’s initial NICU admission lasted 203 days and left her ventilator dependent. Subsequently, she had multiple hospital admissions over the first 18 months of life, mostly for viral respiratory infections.

### SLC25A1 is ubiquitously expressed normal tissues

While Combined D,L-2HGA is a known neurometabolic disorder that affects primarily the central nervous system, the spatial expression pattern of *SLC25A1* in the various organs has not been extensively investigated. To address this, we data mined both GTEx Portal^11^ and The Human Protein Atlas^12^ databases, which revealed ubiquitous expression of *SLC25A1* in a wide variety of tissues and cells, including, but not limited to, cultured fibroblasts, brain, gastrointestinal organs, heart, kidney and liver (Figure 1C). Moreover, segmental analysis of the various compartments of the brain revealed expression of *SLC25A1* in all brain segments, with the highest enrichment localized within the hypothalamus, spinal cord and pituitary gland. At the single cell level, *SLC25A1* is ubiquitously expressed in all neuronal cell types, with the highest enrichment found in oligodendrocytes, inhibitory/excitatory neurons and astrocytes (Supplementary Figure 2)^12^.

### Additional molecular analysis of patient fibroblasts

Given the status of the c.784T>C (p.Cys262Arg) *SLC25A1* variant as a VUS in Patient 893, we further explored its possible pathogenic effects on *SLC25A1* gene expression and function. It is of course possible that a substitution of arginine for cysteine at position 262 impairs CIC stability or function, leading to reduced cellular function. Alternatively, impairment of maturation of the transcribed RNA through altered splicing due to deep intronic splicing pathogenic variants in *SLC25A1* is possible and would be missed due to the lack of capture probes in these non-coding regions in clinical exome sequencing protocols^13,14^. To overcome this limitation, we performed a complementary diagnostic RNA sequencing of the entire transcriptome for FB893 to identify aberrant *de novo* splicing events^15–17^.

Analysis of FB893 fibroblast RNA-seq data using the DROP-FRASER analysis tool that provides a provides a count-based statistical test for aberrant splicing^18,19^ did not identify any atypical spliced transcripts or exon skipping in *SLC25A1* (Supplementary Figure 3) when compared against a cohort of publicly available fibroblast count metrices^20^. Consistent with the WES findings, Exomiser analysis^21^ of the RNA-seq VCF file using random-walk analysis of protein interaction networks, clinical relevance and cross-species phenotype comparisons prioritized the *SLC25A1:* c.784T>C (p.Cys262Arg) homozygous variant (Exomiser Score: 0.987) as the top disease-causing candidate, highly relevant to FB893 clinical presentation (Supplementary Figure 4).

### Combined D,L-2HGA pathophysiology induces a bioenergetic defect in patient derived fibroblasts

Whole cell oxygen consumption studies on patient fibroblasts using a Seahorse XF Bioanalyzer demonstrated impaired cellular bioenergetics in FB893 as evidenced by reduced basal respiration, ATP production, maximal respiration, and spare capacity in FB893 (Figure 3A). Similarly, FB897 fibroblasts derived from the second patient displayed a compromised oxygen consumption profile (Figure 3A). The results collectively support a secondary deficit of mitochondrial oxidative phosphorylation as a consequence of *SLC25A1* mutations. Additional metabolomics analysis is described later.

**Figure 2.**
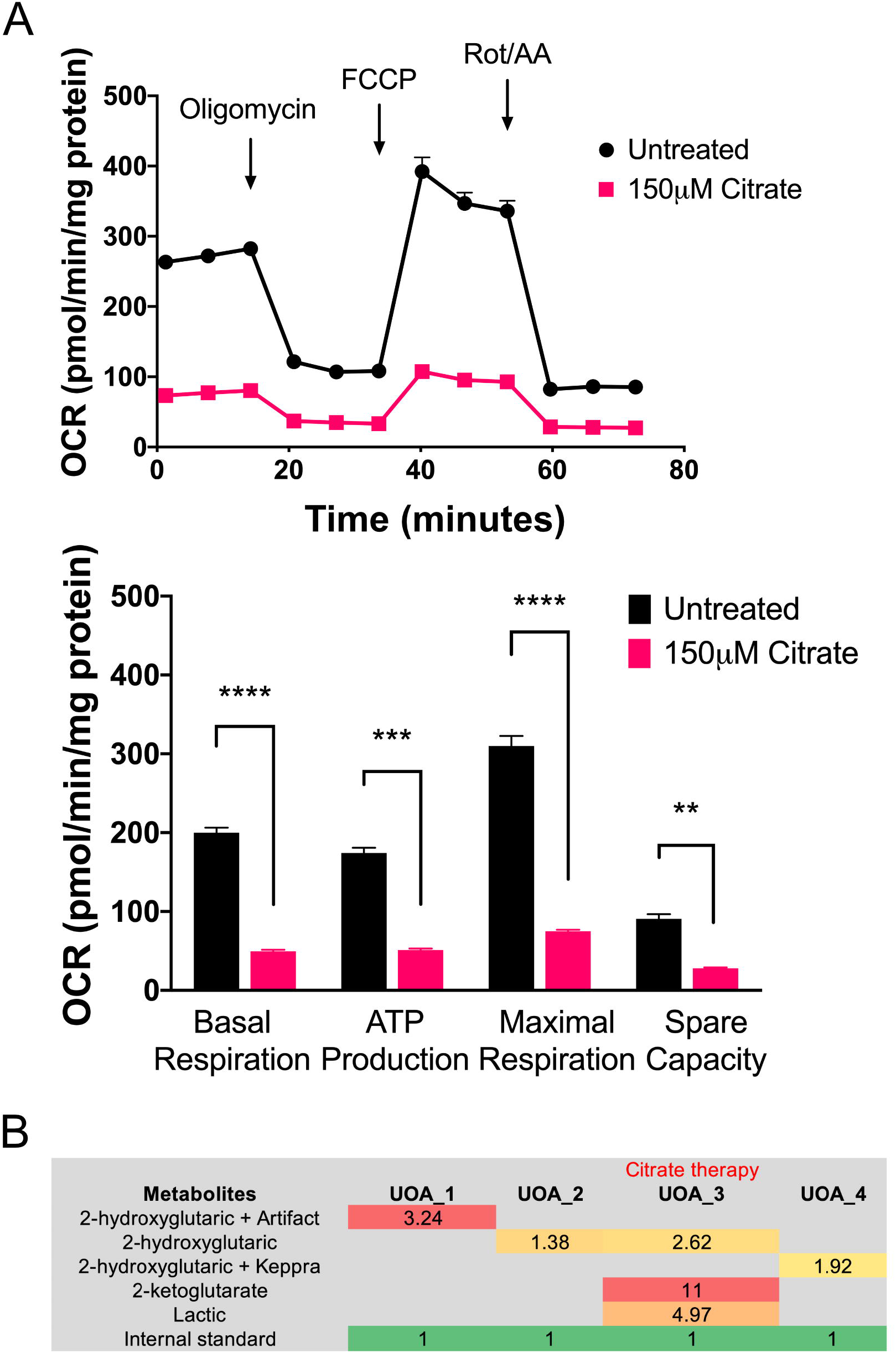
Citrate supplementation of *SLC25A1* deficient patient derived fibroblasts. **(A)** Patient and control fibroblasts were analyzed for whole cell oxygen consumption with a Seahorse bioanalyzer using the company’s MitoStress protocol with and without oxygen consumption. The top graph shows oxygen consumption (pmol/min/mg protein) over time following addition of the indicated compounds. The bottom graph quantitates standard oxygen consumption parameters from these analyses, and shows reduction of all parameters in patient cells compared to control. **Urine organic acid analysis of patient 893 before and after supplementation with citrate. (B)** With citrate supplementation, the patient developed a pattern suggestive of severe metabolic decompensation with excessive secretion of 2-ketoglutarate and 2-hydroxyglutarate. Upon discontinuation of citrate therapy, Patient 893 returned back to clinical baseline conditions. ** p value ≤ 0.01, *** p value ≤ 0.001, **** p value ≤ 0.0001.

**Figure 3.**
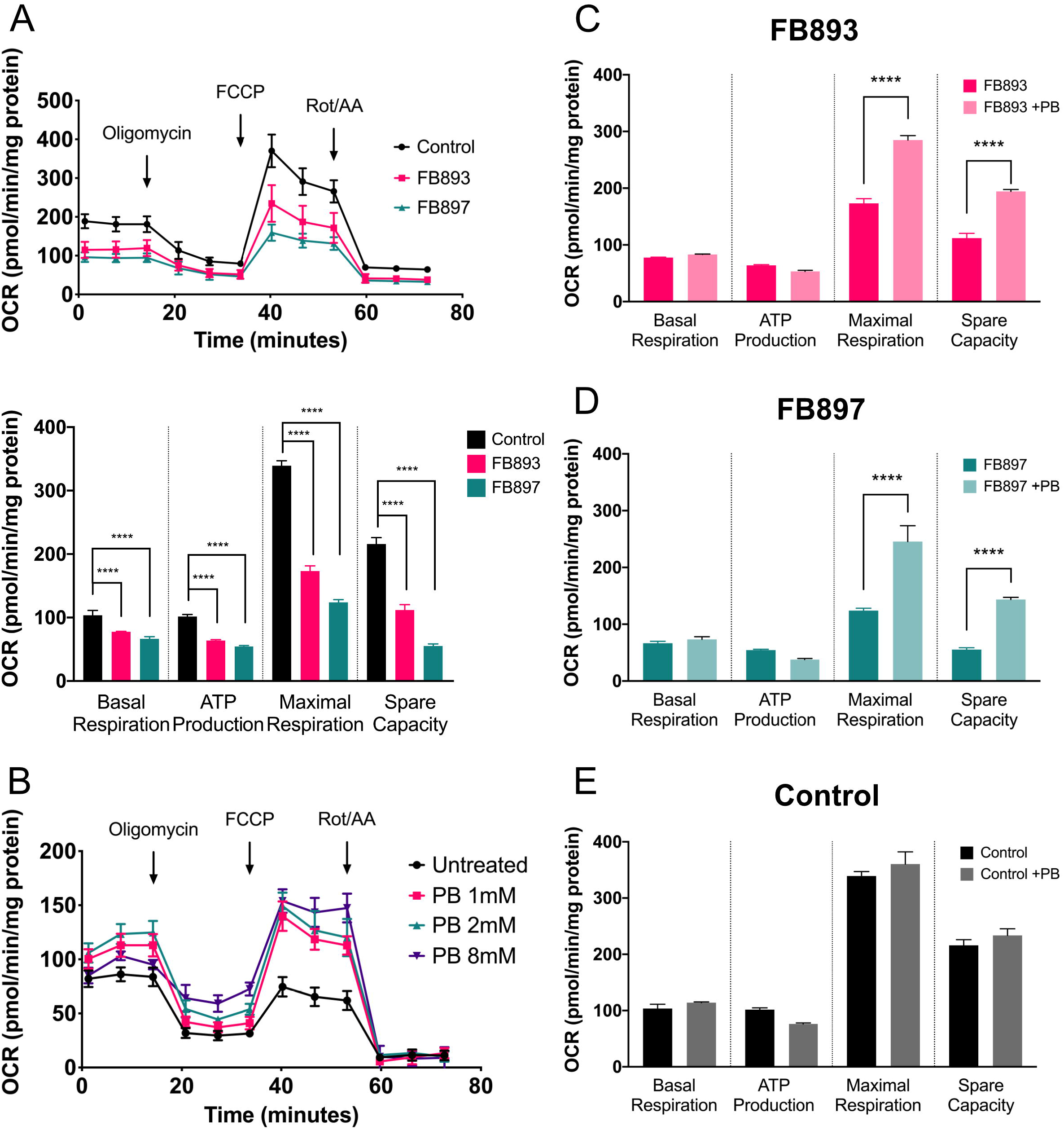
Treatment of patient derived *SLC25A1* deficient fibroblasts with phenylbutyrate improves cellular bioenergetics. **(A)** Assessment of patient derived cells with a Seahorse bioanalyzer confirmed impairment of all parameters including ATP production in cells from both patients. All parameters improved in a dose-dependent fashion with supplementation of 1, 2 or 8 mM phenylbutyrate treatment **(B)** with statistically significant changes in maximal respiration and spare capacity in FB893 **(C)** and FB897 **(D)** but not control cells **(E)**. **** p value ≤ 0.0001.

### Classification of the c.784T>C variant

In lieu of the positive functional findings, we conducted a comprehensive re-evaluation of the c.784T>C (p.Cys262Arg) variant using the ACMG-AMP variant interpretation guidelines^22^ to determine whether the variant qualifies for an upgrade from its initial VUS classification. Population criteria evaluation revealed that his variant is absent in major control population databases, including, gnomAD and ExAC (PM2 criteria)^23,24^. Using UCSC Genome Browser analysis, we verified that the cysteine residue is highly conserved in the vertebrate species, including, human, rhesus, *X.tropicalis* and zebrafish. The high evolutionary conservation indicates that any potential alterations to the amino acid sequence at position 262 is likely to be detrimental (Figure 1E). Missense3D structural analysis^25^ revealed that the wildtype cysteine residue does not form disulfide bonds with any neighboring wildtype amino acids, and that the arginine variant as indicated in this patient’s mutation is not predicted to cause structural damage (Figure 1F). UniProtKB/Swiss-Prot variant analysis^26^ revealed a BLOSUM score of −3, indicating that the wildtype cysteine residue is highly intolerable to amino acid change, and that the arginine substitution causes a physico-chemical property switch from being hydrophobic into hydrophilic. Additionally, the ClinGen predictor REVEL score^27^ of 0.861 for this variant predicts a high likelihood for pathogenicity (PP3_Moderate criteria), and that the functional assay was consistent with a loss of CIC activity (PS3 criteria)^4^. Finally, Patient 893 urine organics profile with an excess secretion of 2-hydroxyglutarate and clinical presentation were highly specific for D,L-2HGA (PP4 criteria). Using these curated information, the c.784T>C variant fulfills PS3, PM2, PP3_Moderate and PP4 criteria, thereby allowing us to re-classify the VUS to likely pathogenic (Supplementary Table 1) which establishes the gene-variant pathogenicity for Patient 893.

### Assessment of citrate therapy in Combined D,L-2HGA

At present, there are no effective treatments for Combined D,L-2HGA and the prognosis for this condition is generally poor. Reduced cytosolic citrate has previously been postulated to be a major D,L-2HGA pathological driver^4^. However, interventional use of citrate remains controversial^4,6^. To further explore citrate supplementation, we cultured FB893 with 150 μM citrate and assessed oxygen consumption with the Seahorse Bioanalzyer. Basal respiration, spare capacity, and ATP production were markedly reduced compared to untreated fibroblasts (Figure 2A). During the course of these *in vitro* experiments, the patient was prescribed citrate supplementation by another care provider, leading to a worsening of clinical symptoms and admission to CHP. Clinical urine organic acid analysis at admission revealed a significant increase in excretion of TCA cycle intermediates including, 2-ketoglutarate and 2-hydroxyglutarate (Figure 2B). Discontinuation of citrate therapy returned the patient to baseline clinical state (Figure 2B). The *in vitro* and clinical findings collectively suggest that citrate therapy is at least contraindicated for Patient 893.

### Treatment with phenylbutyrate improves bioenergetics in Combined D,L-2HGA patient cells

Mutations in *SLC25A1* result in impairment of mitochondrial citrate export, leading to accumulation of citrate that results in buildup of 2-ketoglutarate which eventually gets reduced to the neurotoxic D- and L2-hydroxyglutarate^7,28^. Given the known pathophysiological mechanism of this metabolic disorder, we hypothesized that the reduction of 2-ketoglutarate within the mitochondria would be a viable approach to diminish the level of D- and L2-hydroxyglutarate as well as to normalize mitochondrial homeostasis. Phenylbutyrate is an FDA-approved medication that is used for the management of urea cycle disorder patients through reducing glutamine levels and thus acting as a nitrogen sink^9^. Pharmacokinetically, phenylbutyrate undergoes one cycle of β-oxidation to form phenylacetyl-CoA^8^, which subsequently conjugates to glutamine to form phenylacetylglutamine through the enzyme phenylacetyl-CoA:L-glutamine-N-acetyltransferase, and is eventually excreted from the kidney. Glutamine is made from glutamate, which in turn is made from 2-ketoglutarate (Figure 3A). Thus, phenylbutyrate has the potential to directly reduce the metabolite that leads to D- and L2-hydroxyglutarate production and appears to be a logical candidate for therapy of this disease.

To examine this potential therapy, we treated patient fibroblasts with varying doses of phenylbutyrate to assess the baseline dose dependent efficacy of this drug. In all three doses tested (1, 2 and 8 mM), phenylbutyrate treatment for 72 hrs led to improvement of cellular bioenergetics as evident by increased oxygen consumption with improved maximal respiration and spare capacity (Figure 3B). Both parameters were statistically significantly improved with 8 mM phenylbutyrate treatment in both FB893 and FB897 (Figure 3C, D). Seahorse assessment of phenylbutyrate-treated control fibroblasts demonstrated no discernible changes in oxygen consumption (Figure 3E). Thus, the effect in fibroblast mitochondrial function induced by phenylbutyrate appears to be specific to the Combined D,L-2HGA biochemical defect, rather than a generic off-target effect.

### Treatment of Combined D,L-2HGA fibroblasts and patients by phenylbutyrate

Next, an untargeted metabolomics analysis was performed to assess the change in intracellular metabolites induced by phenylbutyrate treatment. A total of 679 compounds of known identity were detected. Global analysis of the metabolomics data was performed using hierarchal distance clustering based on Euclidean distance metric and revealed two distinct clusters, with the first cluster consisting of controls and phenylbutyrate-treated patient fibroblasts, and the remaining untreated patient fibroblasts forming the second cluster (Figure 4A). In comparison to control fibroblasts, a total of 230 metabolites were found to be differential in the D,L-2HGA fibroblasts based on an adjusted p value ≤ 0.05 (Supplementary Table 2). Consistent with the expected biochemical changes, both FB893 and 897 displayed significant elevation of citrate (29%), 2-ketoglutarate (56%) and 2-hydroxyglutarate (329%) (Figure 4B). The excess levels of 2-ketoglutarate compared to citrate suggests impairment at the 2-ketoglutarate dehydrogenase step, a critical regulatory point in the TCA cycle^28–30^.

**Figure 4.**
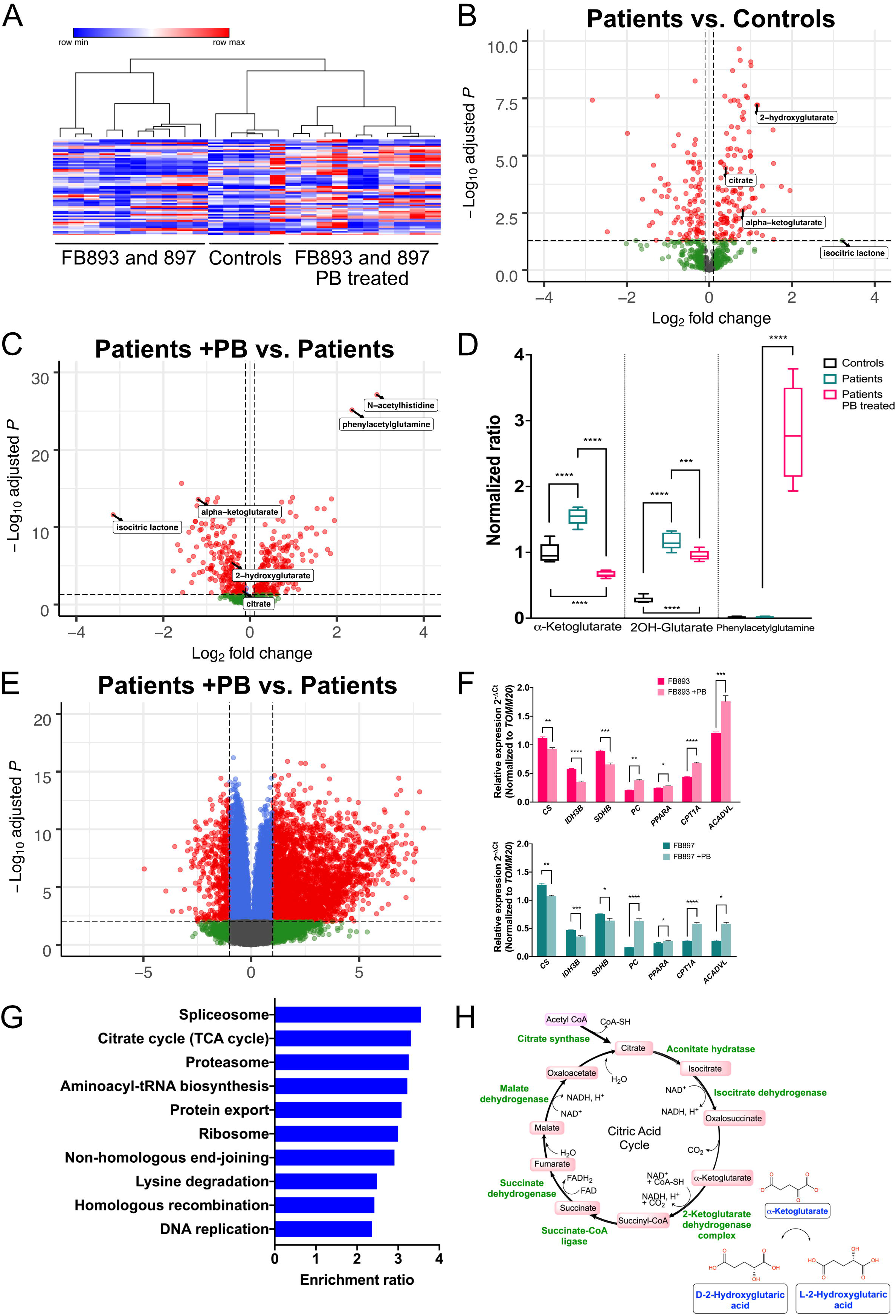
Global metabolomics and RNA-seq analyses demonstrate a positive effects of phenylbutyrate on Combined D,L-2HGA patient derived fibroblasts. **(A)** Hierarchical distance clustering of the metabolomics data demonstrates that phenylbutyrate-treated FB893 and 897 fibroblasts have an improved metabolite profile that closely resembles the control fibroblasts. **(B)** At baseline conditions, both FB893 and 897 displayed significant excess of citrate (29%), 2-ketoglutarate (56%) and 2-hydroxyglutarate (329%) compared to control, consistent with clinical metabolic testing. **(C and D)** Upon phenylbutyrate treatment of FB893 and 897, there was a statistically significant reduction in 2-ketoglutarate (60%) and 2-hydroxyglutarate (20%) that is accompanied by *de novo* formation of phenylacetylglutamine (>16,000 fold). **(E)** RNA-seq analysis of phenylbutyrate-treated FB893 and 897 revealed 7,800 differentially expressed transcripts, with validation of a subset of key genes using qPCR **(F,** FB893 top, and FB897 bottom). **(G)** Pathway analysis of the differentially expressed transcripts showed that phenylbutyrate treatment resulted in differential transcripts including the TCA cycle pathway. **(H)** Transposing the RNA-seq data onto the TCA cycle suggests that the decrease in multiple TCA cycle enzymes at the transcript level could result in reduced carbon flux that would synergize at the metabolite level to augment the reduction of 2-ketoglutarate and D- and L-2-Hydroxyglutaric acids in D,L-2HGA. * p value ≤ 0.05, ** p value ≤ 0.01, *** p value ≤ 0.001, **** p value ≤ 0.0001.

Phenylbutyrate treatment of both FB893 and FB897 resulted in 423 differential metabolites when compared to the untreated state, with a statistically significant reduction in 2-ketoglutarate (60%) and its accompanying analyte 2-hydroxyglutarate (20%) (Figure 4C, D; Supplementary Table 3). Moreover, phenylbutyrate treatment resulted in *de novo* formation of phenylacetylglutamine by >16,000 fold, affirming the presence of an active biochemical reaction between glutamine and phenylacetyl-CoA that is catalyzed by phenylacetyl-CoA:L-glutamine-N-acetyltransferase. These data support the proposition that 2-ketoglutarate depletion strategy in Combined D,L-2HGA via glutamine removal is a logical approach to reduce 2-hydroxyglutarate.

At 18 months of age, Patient 897 had a prolonged hospitalization due to respiratory failure caused by a viral infection that left her ventilator dependent. Patient 897 was subsequently treated with sodium phenylbutyrate at 250 mg/kg/day in 4 divided doses, and within 4 days, she was weaned off her ventilator settings and was soon discharged. After several months on treatment, the family reports improved awareness of surroundings and ability to make eye contact. For the first time she has begun engaging with other individuals and objects. She tolerates up to 4 hours at a time off the ventilator and episodes of respiratory distress requiring additional manual bagging have been eliminated. She has had several viral infections (including COVID-19) since starting sodium phenylbutyrate, none of which have required hospitalization. PAG levels in urine measured clinically by the CLIA approved from near zero (normal) to 9460, then 11622 μg/ml urine, indicative of 2-ketoglutarate from the system. Patient 893 only recently (~3 months) started treatment with Ravicti (glycerol phenylbutyrate) and has not had sufficient time to exhibit a clinical response. However, his urine PAG level has also increased from essentially 0.1 to 149.9 μg/ml urine. Remarkably, his urine 2-hydroxyglutarate was reported to be undetectable through our clinical urine organics analysis (UPMC Children’s Hospital of Pittsburg Clinical Biochemical Genetics Laboratory). Phenylacetate, the derivative analyte from phenylbutyrate, as well as 3OH-Propionate without other propionate analytes were also noted in the urine organics profile (Table 1).

**Table 1.**
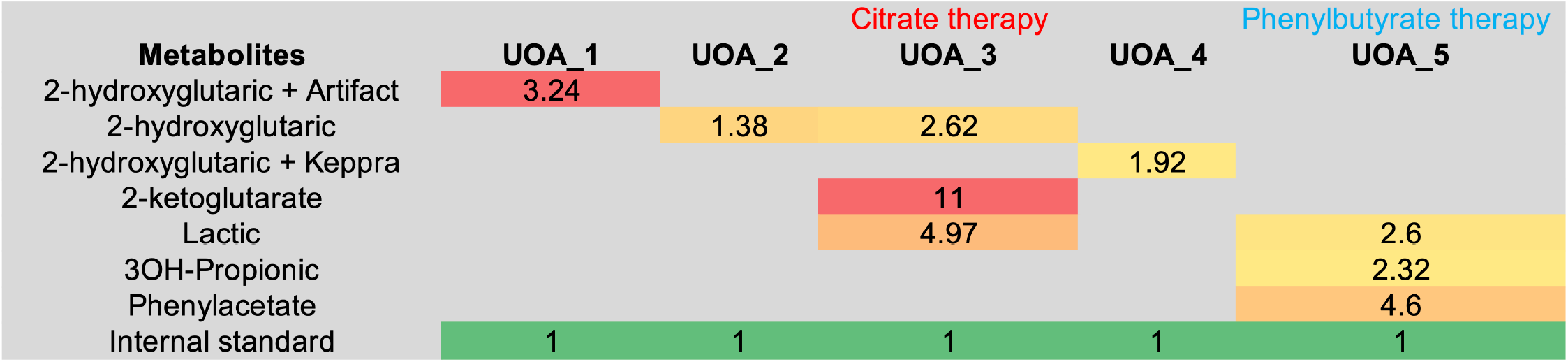
Urine organics profile of Patient 893 following one-month post-phenylbutyrate treatment. GC-MS analysis of Patient 893 urine specimen revealed undetectable 2-hydroxyglutarate. Phenylacetate, a derivative of phenylbutyrate, as well as 3OH-Propionate were observed. The values represent the presence and qualitative ratio of the analyte when compared to the internal standard.

### Differential expression of genes induced by phenylbutyrate treatment of fibroblasts

Given the described inhibitory effect of phenylbutyrate on histone deacetylase^31,32^, we performed RNA-sequencing on the phenylbutyrate treated fibroblasts to identify associated mRNA changes. A total of 7,800 genes were found to be differentially expressed in the phenylbutyrate-treated fibroblasts, with 3,998 and 3,802 upregulated and downregulated respectively (Figure 4E; Supplementary Table 4). To validate the RNA-seq data, we performed qPCR on a relevant subset of genes, and observed a statistically significant downregulation of citrate synthase (*CS*; NM_004077), isocitrate dehydrogenase (NAD^+^) 3 non-catalytic subunit beta (*IDH3B;* NM_006899), succinate dehydrogenase complex iron sulfur subunit B (*SDHB*; NM_003000) and upregulation of pyruvate carboxylase (*PC*; NM_000920), peroxisome proliferator activated receptor alpha (*PPARA;* NM_005036), carnitine palmitoyltransferase 1A (*CPT1A;* NM_001876) and very long chain acyl-CoA dehydrogenase (*ACADVL;* NM_000018) (Figure 4F). Having validated the veracity of the RNA-seq data, we next performed a pathway analysis on the differentially expressed genes using the WebGestalt tool^33^. Using the Over-Representation Analysis approach, we identified a total of 62 categories enriched in our dataset, with the citrate cycle (hsa00020) predicted to be one of the ten highly-enriched pathways based on 70% of curated genes under the TCA cycle category identified to be differentially expressed (21/30 genes) (Figure 4G). By transposing the RNA-seq data onto the TCA cycle, the majority of all key TCA cycle enzyme transcripts were downregulated in response to phenylbutyrate treatment, suggesting that the overall flux of citrate carbon through the TCA cycle is reduce, likely synergizing the metabolic reduction of 2-ketoglutarate and D- and L-2-Hydroxyglutaric in cells, ultimately resulting in improved mitochondrial respiration.

### Compass flux balance analysis through constrained-based optimization predicts improvement in core metabolic pathways upon phenylbutyrate treatment

Compass is a Flux Balance Analysis algorithm that utilizes high dimensional transcriptome changes to deduce metabolic states down to a single-cell resolution level with network-wide comprehensiveness^34^. A key feature of Compass is the ability to allow for global target detection across the entire metabolic network, agnostic of pre-defined metabolic pathway boundaries, and inclusive of underrepresented ancillary pathways that are otherwise critical for cellular function^34,35^. Using Compass, we were able to comprehensively analyze the RNA-seq data against Recon 2 which comprises of 7,440 reactions and 2,626 unique metabolites^36^. In this manner, we identified 1,403 metabolic reactions (Benjamini-Hochberg [BH] adjusted Wilcoxon rank-sum pvalue ≤ 0.05) that are differentially active in phenylbutyrate-treated fibroblasts (Figure 5A, Supplementary Table 5).

**Figure 5.**
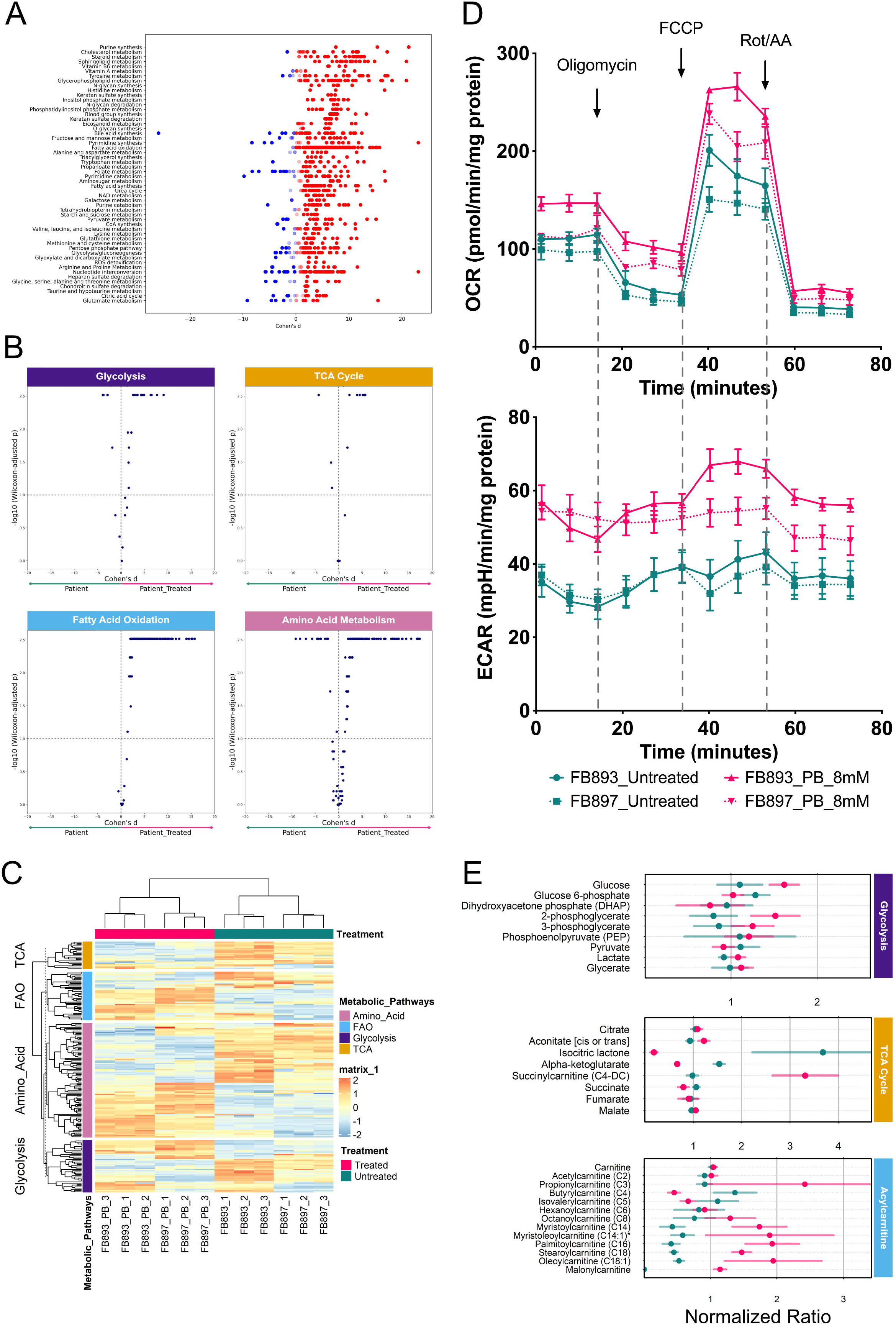
Compass metabolic flux analysis predicts pan-metabolic improvement in Combined D,L-2HGA fibroblasts in response to phenylbutyrate treatment. **(A)** Compass analysis metabolic reactions (dots) are segmented by Recon2 pathways and are highlighted by their corresponding Cohen’s d statistics. **(B)** Compass-score differential activity test predicts an increased activity in the core metabolic pathways. **(C)** Heatmap display of the RNA-seq metabolic genes utilized in Compass analysis. **(D)** Seahorse bioanalyzer analysis revealed an improvement and glycolysis (bottom) in response to phenylbutyrate treatment in addition to the previously identified improvement in mitochondrial respiration (top). **(E)** Metabolomics profiling revealed changes in key metabolites associated with the core metabolic pathways. Each data point represents the average normalized ratio of the individual metabolite measured in phenylbutyrate-treated (pink) and untreated (green) FB893 and 897 fibroblasts, with the error bars denoting the standard deviation. Malonylcarnitine was undetectable in the untreated fibroblasts.

Notably, the Compass algorithm *in silico* prediction indicated that core metabolic pathways including, glycolysis, TCA cycle, fatty acid oxidation and amino acid metabolism were overall improved by phenylbutyrate treatment (Figure 5B, C). Compass predicted that the improvement of glycolysis in the context of enhanced mitochondrial respiration may in part be attributed by downregulation of *PCK2* (logFC: −1.65, pvalue ≤ 0.0001, Supplementary Figure 5, Supplementary Table 4), which has previously been shown to have citrate cataplerosis effects^37,38^, possibly alleviating mitochondrial citrate accumulation, a desirable effect in D,L-2HGA. Fatty acid oxidation appears to be increased, with Compass predicting an increase in carnitine-O-palmitoyltransferase activity, *CPT1A* (logFC: 0.85, p value ≤ 0.0001, Supplementary Table 4), that conjugates carnitine to long chain fatty acyl-CoA, and very long chain acyl-CoA dehydrogrenase *ACADVL*, the first mitochondrial matrix step of fatty acid oxidation (logFC: 0.34, p value ≤ 0.0001, Supplementary Table 4). Finally, Compass predicted multiple amino acid metabolic pathways to be differentially balanced between the untreated and treated state, suggesting a highly regulated fine-tuning of amino acid metabolism.

To validate the Compass predictions, we performed concurrent cellular oxygen consumption (respiration) and proton excretion (glycolysis) analysis through real-time measurements of oxygen consumption rate (OCR) and extracellular acidification rate (ECAR) at intervals of approximately 5-8 minutes using the Seahorse Bioanalyzer. Compared to the untreated state, phenylbutyrate treatment led to enhanced mitochondrial respiration and metabolic fitness as indicated by the responsiveness to FCCP simulated physiological demand for energy consistent with previous results (Figure 3E, F and 5D). Similarly, phenylbutyrate treated fibroblasts displayed a higher ECAR rate, indicating increased lactate excretion into the assay media due to aerobic glycolysis (Figure 5D). Consistent with these findings, metabolomics analysis showed a propensity for higher cellular glycolytic metabolites including, 2-phosphoglycerate, 3-phosphoglycerate and lactate (Figure 5E). Interestingly, we also observed an increase in C3 propionylcarnitine and its derivative C4-DC succinylcarnitine metabolites derived from propiogenic valine and isoleucine amino acids (Figure 5E), suggesting an analeprotic effect on the TCA cycle (Figure 5E). Finally, the higher levels of C14-18 acylcarnitine species is supportive for β-oxidation of very long chain fatty acids and the restoration of malonylcarnitine levels, a surrogate for malonyl-CoA that is the precursor for *de novo* fatty acid synthesis, provides an indirect evidence of enhanced citrate cataplerosis (Figure 5E).

## Discussion

Combined D,L-2HGA is an ultra-rare autosomal recessive neurometabolic disorder that is characterized by severe neonatal-onset encephalopathy, intractable seizures, muscular weakness, respiratory distress and a lack of psychomotor developmental milestones, eventually resulting in early death^1,4–6^. No effective therapy is known.

2-ketoglutarate is a critical intermediate of the TCA cycle, and also plays a vital role in a wide range of physiological processes, including, regulation of catabolic and anabolic TCA cycle metabolites that in turn regulates ATP synthesis, production of NAD^+^/NADH reducing equivalents and amino acid synthesis^28,39,40^. In Combined D,L-2HGA, excess citrate causes accumulation of 2-ketoglutarate, with subsequent conversion into both D- and L-2 hydroxyglutaric acids via the enzymatic activity of hydroxyacid-oxoacid transhydrogenase and L-malate dehydrogenase respectively^1,4,6^. D-2 hydroxyglutaric acid is thought to induce excitotoxic damage to the neurons by activating the N-methyl-d-aspartate (NMDA) receptors, perturbing intracellular calcium homeostasis, oxidative stress and reducing mitochondrial complex V activity^41^. Hence, targeting 2-ketoglutarate and enhancing its reduction represents a compelling therapeutic option to both ameliorate the disequilibrium of the TCA cycle reduce the accumulation of neurotoxic metabolites in this condition. Using a multiomics approach, we have demonstrated that treatment with phenylbutyrate, an FDA-approved medication for urea cycle disorder patients, significantly improved the metabolic state of patient derived fibroblasts with reduced accumulation of 2-ketoglutarate and its toxic derivative 2-hydroxyglutaric acids and a subsequent improvement in cellular bioenergetics. Using a Flux Balance Analysis approach, we showed that phenylbutyrate treatment conferred a pan-metabolic improvement in the core metabolic pathways. Finally, we obtained approval for the use of phenylbutyrate or glycerol phenylbutyrate on two patients with significant neurologic and systemic symptoms. Increased excretion of PAG in both patients confirms that this effect is active *in vivo*. Moreover, the first patient treated showed an improvement in clinical symptoms with no reported adverse events, identifying this drug as a potential therapy for the disorder.

The pathophysiology of Combined D,L-2HGA is felt to be twofold, with the reciprocal depletion of cytosolic citrate induced by the defect proposed to initiate a cascade of diminished substrates, including oxaloacetate and nicotinamide adenine dinucleotide phosphate (NADPH). In addition, acetyl-CoA derived malonyl-CoA is required for fatty acid, sterol and plasmalogen biosynthesis^42–45^. Consequently, dysregulated lipid metabolism is thought to elicit a multifaceted neurological effect^45–47^ that underlies the majority of the neurological dysfunction in Combined D,L-2HGA^4^. Phenylbutyrate is not likely to affect this metabolic abnormality in patients. Thus, the possibility of concurrent citrate supplementation must be considered. Citrate therapy has previously been reported to stabilize the severity and frequency of seizures in a Combined D,L-2HGA patient, presumably due to repletion of the cytoplasmic citrate pool^4^. However, the treatment paradoxically increased patient urinary excretion of TCA cycle intermediates including malate and succinate in response to the therapy, possibly reflecting an increase in mitochondrial stress. Our Patient 893 experienced an acute metabolic decompensation associated with citrate supplementation, with metabolite evidence of mitochondrial stress that resolved upon discontinuation of citrate and worsening fibroblast bioenergetics. The latter result is in agreement with a previous report that treatment of *SLC25A1*-deficient fibroblasts with 4 mM citrate led to reduced cellular spare respiratory capacity and lack of sustained improvement in the patient^6^. Thus, the merits of citrate therapy for this disorder remain questionable.

The reduction of 2-ketoglutarate and D,L-2- Hydroxyglutarate in response to phenylbutyrate may not be the sole contributing factor in improving bioenergetics in D,L-2HGA fibroblasts. Our RNA-seq and flux balance analysis via constrained-based optimization revealed transcriptional changes in response to phenylbutyrate treatment that appears to exhibit a pan-metabolic improvement in the core metabolic pathways. A key observation in our transcriptome analysis is the downregulation of *PCK2*, a mitochondrial phosphoenolpyruvate carboxykinase that is known to play a central role in regulating and fine-tuning between glycolysis, TCA cycle, and gluconeogenesis during metabolic adaption^37,48,49^. Downregulation of *PCK2* has previously been shown to promote citrate cataplerosis with citrate carbon outflow from the mitochondrial into the cytosol, leading to increased cytosolic acetyl-CoA-fatty acid biogenesis^37^. In agreement with the citrate cataplerosis effect, phenylbutyrate-treated patient fibroblasts resulted in restoration of malonylcarnitine, indicating an elevation of malonyl-CoA that can support biosynthesis of fatty acids in the cytoplasm^42,50^. While downregulation of *PCK2* and mitochondrial citrate results in reduced oxidative phosphorylation in B16 tumor-repopulating cells^37^, the paradoxical improvement in oxidative phosphorylation in phenylbutyrate-treated Combined D,L-2HGA fibroblasts further underscores the critical aspect of normalizing mitochondrial citrate to physiological levels in improving bioenergetics in CIC deficient fibroblasts.

In conclusion, we have identified phenylbutyrate, the active ingredient of triphenylbutyrylglycerol (commercially known as Ravicti) as potential therapeutic agent for Combined D,L-2HGA. Phenylbutyrate has the potential to directly reduce the accumulation of the toxic metabolite as well as induce important secondary changes in a variety of metabolic pathways. Also, we cautioned against the use of citrate as an approach to compensate for its cytoplasmic depletion. Finally, the improvement in cellular bioenergetics in patient derived fibroblasts in conjunction with decrease in mitochondrial citrate, 2-ketoglutarate, and its derivative 2-hydroxyglutarate derivatives to physiological levels underscores the metabolic relevance of these compounds in improving pathophysiology in this disease.

## Acknowledgements

YLP was an ABMGG Clinical Biochemical Genetics fellow supported by a pilot project grant 20(GG014929-16) from North American Mitochondrial Disease Consortium under the Rare Diseases Clinical Research Network and a Rangos Advisory Committee fellowship grant from UPMC Children’s Hospital of Pittsburgh. YLP is currently an ABMGG Laboratory Genetics and Genomics fellow supported by the Icahn School of Medicine at Mount Sinai Clinical Genetics Laboratory Training Program. JV is the Cleveland Family Endowed Chair in Pediatric Research with support in part from NIH grant R01DK109907. We acknowledge our patients and families for their participation in this study, Dr. Johan Van Hove for advice on the NAMDC IRB application, the University of Pittsburgh Rangos Metabolic Core services, Center for Research Computing and the clinical diagnostic services provided by the General Genetics Clinic and Clinical Biochemical Genetics Laboratory.

## References

1. Nota, B., Struys, E.A., Pop, A., Jansen, E.E., Fernandez Ojeda, M.R., Kanhai, W.A., Kranendijk, M., van Dooren, S.J., Bevova, M.R., Sistermans, E.A., et al. (2013). Deficiency in SLC25A1, encoding the mitochondrial citrate carrier, causes combined D-2- and L-2-hydroxyglutaric aciduria. Am J Hum Genet 92, 627–631. 10.1016/j.ajhg.2013.03.009.

2. Chaouch, A., Porcelli, V., Cox, D., Edvardson, S., Scarcia, P., De Grassi, A., Pierri, C.L., Cossins, J., Laval, S.H., Griffin, H., et al. (2014). Mutations in the Mitochondrial Citrate Carrier SLC25A1 are Associated with Impaired Neuromuscular Transmission. J Neuromuscul Dis 1, 75–90. 10.3233/JND-140021.

3. Pop, A., Williams, M., Struys, E.A., Monne, M., Jansen, E.E.W., De Grassi, A., Kanhai, W.A., Scarcia, P., Ojeda, M.R.F., Porcelli, V., et al. (2018). An overview of combined D-2- and L-2-hydroxyglutaric aciduria: functional analysis of CIC variants. J Inherit Metab Dis 41, 169–180. 10.1007/s10545-017-0106-7.

4. Muhlhausen, C., Salomons, G.S., Lukacs, Z., Struys, E.A., van der Knaap, M.S., Ullrich, K., and Santer, R. (2014). Combined D2-/L2-hydroxyglutaric aciduria (SLC25A1 deficiency): clinical course and effects of citrate treatment. J Inherit Metab Dis 37, 775–781. 10.1007/s10545-014-9702-y.

5. Prasun, P., Young, S., Salomons, G., Werneke, A., Jiang, Y.H., Struys, E., Paige, M., Avantaggiati, M.L., and McDonald, M. (2015). Expanding the Clinical Spectrum of Mitochondrial Citrate Carrier (SLC25A1) Deficiency: Facial Dysmorphism in Siblings with Epileptic Encephalopathy and Combined D,L-2- Hydroxyglutaric Aciduria. JIMD Rep 19, 111–115. 10.1007/8904_2014_378.

6. Smith, A., McBride, S., Marcadier, J.L., Michaud, J., Al-Dirbashi, O.Y., Schwartzentruber, J., Beaulieu, C.L., Katz, S.L., Consortium, F.C., Majewski, J., et al. (2016). Severe Neonatal Presentation of Mitochondrial Citrate Carrier (SLC25A1) Deficiency. JIMD Rep 30, 73–79. 10.1007/8904_2016_536.

7. Catalina-Rodriguez, O., Kolukula, V.K., Tomita, Y., Preet, A., Palmieri, F., Wellstein, A., Byers, S., Giaccia, A.J., Glasgow, E., Albanese, C., and Avantaggiati, M.L. (2012). The mitochondrial citrate transporter, CIC, is essential for mitochondrial homeostasis. Oncotarget 3, 1220–1235. 10.18632/oncotarget.714.

8. Kormanik, K., Kang, H., Cuebas, D., Vockley, J., and Mohsen, A.W. (2012). Evidence for involvement of medium chain acyl-CoA dehydrogenase in the metabolism of phenylbutyrate. Mol Genet Metab 107, 684–689. 10.1016/j.ymgme.2012.10.009.

9. Berry, S.A., Longo, N., Diaz, G.A., McCandless, S.E., Smith, W.E., Harding, C.O., Zori, R., Ficicioglu, C., Lichter-Konecki, U., Robinson, B., and Vockley, J. (2017). Safety and efficacy of glycerol phenylbutyrate for management of urea cycle disorders in patients aged 2months to 2years. Mol Genet Metab 122, 46–53. 10.1016/j.ymgme.2017.09.002.

10. Dobrowolski, S.F., Alodaib, A., Karunanidhi, A., Basu, S., Holecko, M., Lichter-Konecki, U., Pappan, K.L., and Vockley, J. (2020). Clinical, biochemical, mitochondrial, and metabolomic aspects of methylmalonate semialdehyde dehydrogenase deficiency: Report of a fifth case. Mol Genet Metab 129, 272–277. 10.1016/j.ymgme.2020.01.005.

11. Consortium, G.T. (2013). The Genotype-Tissue Expression (GTEx) project. Nat Genet 45, 580–585. 10.1038/ng.2653.

12. Uhlen, M., Fagerberg, L., Hallstrom, B.M., Lindskog, C., Oksvold, P., Mardinoglu, A., Sivertsson, A., Kampf, C., Sjostedt, E., Asplund, A., et al. (2015). Proteomics. Tissue-based map of the human proteome. Science 347, 1260419. 10.1126/science.1260419.

13. Sun, Y., Ruivenkamp, C.A., Hoffer, M.J., Vrijenhoek, T., Kriek, M., van Asperen, C.J., den Dunnen, J.T., and Santen, G.W. (2015). Next-generation diagnostics: gene panel, exome, or whole genome? Hum Mutat 36, 648–655. 10.1002/humu.22783.

14. Vaz-Drago, R., Custodio, N., and Carmo-Fonseca, M. (2017). Deep intronic mutations and human disease. Hum Genet 136, 1093–1111. 10.1007/s00439-017-1809-4.

15. Cummings, B.B., Marshall, J.L., Tukiainen, T., Lek, M., Donkervoort, S., Foley, A.R., Bolduc, V., Waddell, L.B., Sandaradura, S.A., O’Grady, G.L., et al. (2017). Improving genetic diagnosis in Mendelian disease with transcriptome sequencing. Sci Transl Med 9. 10.1126/scitranslmed.aal5209.

16. Kremer, L.S., Bader, D.M., Mertes, C., Kopajtich, R., Pichler, G., Iuso, A., Haack, T.B., Graf, E., Schwarzmayr, T., Terrile, C., et al. (2017). Genetic diagnosis of Mendelian disorders via RNA sequencing. Nat Commun 8, 15824. 10.1038/ncomms15824.

17. Fresard, L., Smail, C., Ferraro, N.M., Teran, N.A., Li, X., Smith, K.S., Bonner, D., Kernohan, K.D., Marwaha, S., Zappala, Z., et al. (2019). Identification of rare-disease genes using blood transcriptome sequencing and large control cohorts. Nat Med 25, 911–919. 10.1038/s41591-019-0457-8.

18. Yepez, V.A., Mertes, C., Muller, M.F., Klaproth-Andrade, D., Wachutka, L., Fresard, L., Gusic, M., Scheller, I.F., Goldberg, P.F., Prokisch, H., and Gagneur, J. (2021). Detection of aberrant gene expression events in RNA sequencing data. Nat Protoc 16, 1276–1296. 10.1038/s41596-020-00462-5.

19. Mertes, C., Scheller, I.F., Yepez, V.A., Celik, M.H., Liang, Y., Kremer, L.S., Gusic, M., Prokisch, H., and Gagneur, J. (2021). Detection of aberrant splicing events in RNA-seq data using FRASER. Nat Commun 12, 529. 10.1038/s41467-020-20573-7.

20. Yepez, V.A., Gusic, M., Kopajtich, R., Mertes, C., Smith, N.H., Alston, C.L., Ban, R., Beblo, S., Berutti, R., Blessing, H., et al. (2022). Clinical implementation of RNA sequencing for Mendelian disease diagnostics. Genome Med 14, 38. 10.1186/s13073-022-01019-9.

21. Smedley, D., Jacobsen, J.O., Jager, M., Kohler, S., Holtgrewe, M., Schubach, M., Siragusa, E., Zemojtel, T., Buske, O.J., Washington, N.L., et al. (2015). Next-generation diagnostics and disease-gene discovery with the Exomiser. Nat Protoc 10, 2004–2015. 10.1038/nprot.2015.124.

22. Richards, S., Aziz, N., Bale, S., Bick, D., Das, S., Gastier-Foster, J., Grody, W.W., Hegde, M., Lyon, E., Spector, E., et al. (2015). Standards and guidelines for the interpretation of sequence variants: a joint consensus recommendation of the American College of Medical Genetics and Genomics and the Association for Molecular Pathology. Genet Med 17, 405–424. 10.1038/gim.2015.30.

23. Karczewski, K.J., Francioli, L.C., Tiao, G., Cummings, B.B., Alfoldi, J., Wang, Q., Collins, R.L., Laricchia, K.M., Ganna, A., Birnbaum, D.P., et al. (2020). The mutational constraint spectrum quantified from variation in 141,456 humans. Nature 581, 434–443. 10.1038/s41586-020-2308-7.

24. Lek, M., Karczewski, K.J., Minikel, E.V., Samocha, K.E., Banks, E., Fennell, T., O’Donnell-Luria, A.H., Ware, J.S., Hill, A.J., Cummings, B.B., et al. (2016). Analysis of protein-coding genetic variation in 60,706 humans. Nature 536, 285–291. 10.1038/nature19057.

25. Ittisoponpisan, S., Islam, S.A., Khanna, T., Alhuzimi, E., David, A., and Sternberg, M.J.E. (2019). Can Predicted Protein 3D Structures Provide Reliable Insights into whether Missense Variants Are Disease Associated? J Mol Biol 431, 2197–2212. 10.1016/j.jmb.2019.04.009.

26. UniProt, C. (2021). UniProt: the universal protein knowledgebase in 2021. Nucleic Acids Res 49,D480-D489. 10.1093/nar/gkaa1100.

27. Ioannidis, N.M., Rothstein, J.H., Pejaver, V., Middha, S., McDonnell, S.K., Baheti, S., Musolf, A., Li, Q., Holzinger, E., Karyadi, D., et al. (2016). REVEL: An Ensemble Method for Predicting the Pathogenicity of Rare Missense Variants. Am J Hum Genet 99, 877–885. 10.1016/j.ajhg.2016.08.016.

28. Martinez-Reyes, I., and Chandel, N.S. (2020). Mitochondrial TCA cycle metabolites control physiology and disease. Nat Commun 11, 102. 10.1038/s41467-019-13668-3.

29. Qi, F., Pradhan, R.K., Dash, R.K., and Beard, D.A. (2011). Detailed kinetics and regulation of mammalian 2-oxoglutarate dehydrogenase. BMC Biochem 12, 53. 10.1186/1471-2091-12-53.

30. Vatrinet, R., Leone, G., De Luise, M., Girolimetti, G., Vidone, M., Gasparre, G., and Porcelli, A.M. (2017). The alpha-ketoglutarate dehydrogenase complex in cancer metabolic plasticity. Cancer Metab 5, 3. 10.1186/s40170-017-0165-0.

31. Kusaczuk, M., Kretowski, R., Bartoszewicz, M., and Cechowska-Pasko, M. (2016). Phenylbutyrate-a pan-HDAC inhibitor-suppresses proliferation of glioblastoma LN-229 cell line. Tumour Biol 37, 931–942. 10.1007/s13277-015-3781-8.

32. Sandner, G., Host, L., Angst, M.J., Guiberteau, T., Guignard, B., and Zwiller, J. (2011). The HDAC Inhibitor Phenylbutyrate Reverses Effects of Neonatal Ventral Hippocampal Lesion in Rats. Front Psychiatry 1, 153. 10.3389/fpsyt.2010.00153.

33. Liao, Y., Wang, J., Jaehnig, E.J., Shi, Z., and Zhang, B. (2019). WebGestalt 2019: gene set analysis toolkit with revamped UIs and APIs. Nucleic Acids Res 47, W199–W205. 10.1093/nar/gkz401.

34. Wagner, A., Wang, C., Fessler, J., DeTomaso, D., Avila-Pacheco, J., Kaminski, J., Zaghouani, S., Christian, E., Thakore, P., Schellhaass, B., et al. (2021). Metabolic modeling of single Th17 cells reveals regulators of autoimmunity. Cell 184, 4168–4185 e4121. 10.1016/j.cell.2021.05.045.

35. Puleston, D.J., Villa, M., and Pearce, E.L. (2017). Ancillary Activity: Beyond Core Metabolism in Immune Cells. Cell Metab 26, 131–141. 10.1016/j.cmet.2017.06.019.

36. Thiele, I., Swainston, N., Fleming, R.M., Hoppe, A., Sahoo, S., Aurich, M.K., Haraldsdottir, H., Mo, M.L., Rolfsson, O., Stobbe, M.D., et al. (2013). A community-driven global reconstruction of human metabolism. Nat Biotechnol 31, 419–425. 10.1038/nbt.2488.

37. Luo, S., Li, Y., Ma, R., Liu, J., Xu, P., Zhang, H., Tang, K., Ma, J., Liu, N., Zhang, Y., et al. (2017). Downregulation of PCK2 remodels tricarboxylic acid cycle in tumor-repopulating cells of melanoma. Oncogene 36, 3609–3617. 10.1038/onc.2016.520.

38. Zhao, J., Li, J., Fan, T.W.M., and Hou, S.X. (2017). Glycolytic reprogramming through PCK2 regulates tumor initiation of prostate cancer cells. Oncotarget 8, 83602–83618. 10.18632/oncotarget.18787.

39. Yuan, Y., Zhu, C., Wang, Y., Sun, J., Feng, J., Ma, Z., Li, P., Peng, W., Yin, C., Xu, G., et al. (2022). alpha-Ketoglutaric acid ameliorates hyperglycemia in diabetes by inhibiting hepatic gluconeogenesis via serpina1e signaling. Sci Adv 8, eabn2879. 10.1126/sciadv.abn2879.

40. Zdzisinska, B., Zurek, A., and Kandefer-Szerszen, M. (2017). Alpha-Ketoglutarate as a Molecule with Pleiotropic Activity: Well-Known and Novel Possibilities of Therapeutic Use. Arch Immunol Ther Exp (Warsz) 65, 21–36. 10.1007/s00005-016-0406-x.

41. Kolker, S., Pawlak, V., Ahlemeyer, B., Okun, J.G., Horster, F., Mayatepek, E., Krieglstein, J., Hoffmann, G.F., and Kohr, G. (2002). NMDA receptor activation and respiratory chain complex V inhibition contribute to neurodegeneration in d-2-hydroxyglutaric aciduria. Eur J Neurosci 16, 21–28. 10.1046/j.1460-9568.2002.02055.x.

42. Foster, D.W. (2012). Malonyl-CoA: the regulator of fatty acid synthesis and oxidation. J Clin Invest 122, 1958–1959. 10.1172/jci63967.

43. Liu, D., Xiao, Y., Evans, B.S., and Zhang, F. (2015). Negative feedback regulation of fatty acid production based on a malonyl-CoA sensor-actuator. ACS Synth Biol 4, 132–140. 10.1021/sb400158w.

44. Wakil, S.J., and Abu-Elheiga, L.A. (2009). Fatty acid metabolism: target for metabolic syndrome. J Lipid Res 50 Suppl, S138–143. 10.1194/jlr.R800079-JLR200.

45. Tracey, T.J., Steyn, F.J., Wolvetang, E.J., and Ngo, S.T. (2018). Neuronal Lipid Metabolism: Multiple Pathways Driving Functional Outcomes in Health and Disease. Front Mol Neurosci 11, 10. 10.3389/fnmol.2018.00010.

46. Hussain, G., Wang, J., Rasul, A., Anwar, H., Imran, A., Qasim, M., Zafar, S., Kamran, S.K.S., Razzaq, A., Aziz, N., et al. (2019). Role of cholesterol and sphingolipids in brain development and neurological diseases. Lipids Health Dis 18, 26. 10.1186/s12944-019-0965-z.

47. Martin, M.G., Pfrieger, F., and Dotti, C.G. (2014). Cholesterol in brain disease: sometimes determinant and frequently implicated. EMBO Rep 15, 1036–1052. 10.15252/embr.201439225.

48. Vincent, E.E., Sergushichev, A., Griss, T., Gingras, M.C., Samborska, B., Ntimbane, T., Coelho, P.P., Blagih, J., Raissi, T.C., Choiniere, L., et al. (2015). Mitochondrial Phosphoenolpyruvate Carboxykinase Regulates Metabolic Adaptation and Enables Glucose-Independent Tumor Growth. Mol Cell 60, 195–207. 10.1016/j.molcel.2015.08.013.

49. Montal, E.D., Dewi, R., Bhalla, K., Ou, L., Hwang, B.J., Ropell, A.E., Gordon, C., Liu, W.J., DeBerardinis, R.J., Sudderth, J., et al. (2015). PEPCK Coordinates the Regulation of Central Carbon Metabolism to Promote Cancer Cell Growth. Mol Cell 60, 571–583. 10.1016/j.molcel.2015.09.025.

50. McGarry, J.D., Mannaerts, G.P., and Foster, D.W. (1977). A possible role for malonyl-CoA in the regulation of hepatic fatty acid oxidation and ketogenesis. J Clin Invest 60, 265–270. 10.1172/JCI108764.

