## Supplementary Data for "A multiomics approach to understanding pathology of Combined D,L-2- Hydroxyglutaric Aciduria and phenylbutyrate as potential treatment"

Detailed Materials and Methods

Supplementary Figure 1. Magnetic Resonance Imaging showed severe white matter loss.
Supplementary Figure 2. *SLC25A1* is ubiquitously expressed in all neuronal cells.
Supplementary Figure 3. DROP-FRASER and Sashimi plot analysis revealed no evidence of aberrant spliced *SLC25A1* transcripts.
Supplementary Figure 4. Exomiser analysis of FB893 RNA-seq prioritizes *SLC25A1* as the top disease gene candidate.
Supplementary Figure 5. RNA-seq showed a reduction of *PCK2* in phenylbutyrate-treated fibroblasts.

Supplementary Table 1. *SLC25A1* variant information and ACMG-AMP classification.
Supplementary Table 2. Differential metabolites in Combined D,L-2HGA fibroblasts at baseline conditions.
Supplementary Table 3. Differential metabolites in phenylbutyrate-treated Combined D,L-2HGA fibroblasts.
Supplementary Table 4. Differentially expressed genes in phenylbutyrate-treated Combined D,L-2HGA fibroblasts.
Supplementary Table 5. Compass in-silico prediction of differential metabolic flux in phenylbutyrate-treated Combined D,L-2HGA fibroblasts.
Supplementary Table 6. List of qPCR primers used in this study.

Supplementary References

### Detailed Materials and Methods

#### *Assessment of whole cell mitochondrial respiration through oximetry*

Fibroblasts were seeded in 96-well Seahorse tissue culture microplates (Agilent #101085-004) at a density of 40,000 cells per well and treated with phenylbutyrate at a final concentration of 8 mM in 100 µL complete DMEM media for 72 hrs in a humidified 37 °C, 5% $CO_2$ incubator. For each cell line, 8 replicate wells were used per experimental condition. The oxygen consumption rate (OCR) was measured with the use of a Seahorse XF Cell Mito Stress Test Kit (Agilent #103015-100) according to the manufacturer's protocol. In brief, one day prior to the assessment, the XFe96 FluxPak (Agilent #102416-100) was hydrated overnight using the Seahorse XF Calibrant Solution (Agilent #100840-000) in a humidified 37 °C non-$CO_2$ incubator. On the day of the assessment, the seeded microplate was washed twice with Seahorse XF DMEM media (Agilent #103575-100), followed by a 1 hr incubation in in the same media supplemented with 10 mM glucose (Agilent #103577-100) and 2 mM glutamine (Agilent #103579-100) in a humidified 37 °C non-$CO_2$ incubator. Oligomycin, FCCP and Rot/AA compounds supplied in the Mito Stress Test Kit were prepared using Seahorse XF DMEM media and used at a final well concentration of 1.5 µM, 1.0 µM and 0.5 µM respectively. Fibroblast OCR was measured on a Seahorse XFe96 Analyzer (Agilent Technologies Inc) using the default XF Cell Mito Stress protocol on the Seahorse Wave Controller Software v2.6.1. Following completion of the Seahorse assessment, 20 µL of RIPA buffer (ThermoFisher Scientific #89900) was added to the cells and 5 µL of lysate was used for protein quantification using a DC Protein Assay Kit II (Bio-Rad #5000112). The OCR reading of each well was normalized to its protein concentration.

#### *RNA-sequencing and quantitative PCR*

Three replicates of each cell line (experimental, control) per experimental condition were grown to 80% confluence in T175 flasks. Following 72 hrs of phenylbutyrate treatment in complete DMEM media, the cells were washed once with phosphate buffered saline (PBS; ThermoFisher Scientific #10010023) and trypsinized using 0.25% trypsin (Corning Life Sciences #25-053-CI). The collected cell pellets were washed once with PBS and processed for total RNA extraction using a Qiagen RNeasy Mini Kit (Qiagen #74104) with on-column DNase digestion (Qiagen #79254). RNA was eluted using nuclease-free water, followed by integrity assessment using a Fragment Analyzer System (Agilent Technologies Inc) prior to storage at -80 °C. Total RNA was submitted to GENEWIZ (South Plainfield, NJ), an Illumina CSPro certified laboratory, for Illumina HiSeq 2X150 bp sequencing using a PolyA selection approach with ~50 million reads per sample. The resulting FASTQ files were first aligned and mapped to the hg38 genome using Rsubread, followed by gene annotation and counting using featureCounts Bioconductor packages in R[1,2]. Statistical analysis was performed using limma-voom package, and genes with a positive B-statistics value were considered statistically significant[3,4]. Pathway analysis was performed using the WebGestalt tool[5,6].

The VCF file from the RNA-seq was generated in accordance to GATK RNAseq short variant discovery (SNPs + Indels) best practices workflow[7], followed by Exomiser analysis to prioritize genes and variants for novel disease-gene discovery or differential diagnosis of Mendelian disease[8]. Aberrant splicing event analysis was performed using the DROP-FRASER tool[9,10], and the RNA-seq data was mapped to the GENCODE hg19 Release 34[11] using the STAR aligner twoPassMode Basic option[12]. External count matrices for 154 fibroblast samples which were previously generated through DROP using the GENCODE hg19 Release 34 were retrieved from Zenodo (doi: 10.5281/zenodo.4646823) for comparison purposes[13]. Metabolic flux prediction and analysis of the RNA-seq data was performed using the Compass algorithm[14]. All analyses were performed using the High Performance Computing resources at the University of Pittsburgh Center for Research Computing.

For quantitative PCR assessment, 2500ng of total RNA were reverse transcribed into cDNA using a SuperScript IV VILO Master Mix Kit (ThermoFisher Scientific #11756050). Quantitative PCR was carried out using PowerUp SYBR Green Master Mix (ThermoFisher Scientific #A25742) on a Bio-Rad CFX96 Real-Time PCR Instrument, with threshold cycle (Ct) values normalized to *TOMM20* and the relative expression computed using the $2^{-\Delta Ct}$ method. PCR primers are listed in Supplementary Table 6.

***Metabolomics***

Five replicates of fibroblasts per cell line (experimental, control) per experimental condition were grown to 80% confluence in T175 flasks with or without phenylbutyrate treatment in complete DMEM media. Following 72 hrs of phenylbutyrate treatment, the cells were harvested as described earlier and stored at -80 °C prior to submission to Metabolon (Morrisville, NC) for global metabolomics analysis as previously described[15,16]. Per Metabolon, samples were prepared using the automated MicroLab STAR® system from Hamilton Company. Several recovery standards were added prior to extraction for QC purposes. To remove protein and dissociate small molecules bound to protein or trapped in the precipitated protein matrix, proteins were precipitated with methanol under vigorous shaking for 2 min (Glen Mills GenoGrinder 2000) followed by centrifugation. The resulting extract was divided into five fractions: two for analysis by two separate reverse phase (RP)/UPLC-MS/MS methods with positive ion mode electrospray ionization (ESI), one for analysis by RP/UPLC-MS/MS with negative ion mode ESI, one for analysis by HILIC/UPLC-MS/MS with negative ion mode ESI, and one sample was reserved for backup. Samples were placed briefly on a TurboVap® (Zymark) to remove the organic solvent. The sample extracts were stored overnight under nitrogen before preparation for analysis.

All analytical methods utilized a Waters ACQUITY ultra-performance liquid chromatography (UPLC) and a Thermo Scientific Q-Exactive high resolution/accurate mass spectrometer interfaced with a heated electrospray ionization (HESI-II) source and Orbitrap mass analyzer operated at 35,000 mass resolution. The sample extract was dried then reconstituted in solvents compatible to each of the four methods. Each reconstitution solvent contained a series of standards at fixed concentrations to ensure injection and chromatographic consistency. One aliquot was analyzed using acidic positive ion conditions, chromatographically optimized for more hydrophilic compounds. In this method, the extract was gradient eluted from a C18 column (Waters UPLC BEH C18-2.1x100 mm, 1.7 µm) using water and methanol, containing 0.05% perfluoropentanoic acid (PFPA) and 0.1% formic acid (FA). Another aliquot was also analyzed using acidic positive ion conditions; however it was chromatographically optimized for more hydrophobic compounds. In this method, the extract was gradient eluted from the same afore mentioned C18 column using methanol, acetonitrile, water, 0.05% PFPA and 0.01% FA and was operated at an overall higher organic content. Another aliquot was analyzed using basic negative ion optimized conditions using a separate dedicated C18 column. The basic extracts were gradient eluted from the column using methanol and water, however with 6.5 mM Ammonium Bicarbonate at pH 8. The fourth aliquot was analyzed via negative ionization following elution from a HILIC column (Waters UPLC BEH Amide 2.1x150 mm, 1.7 µm) using a gradient consisting of water and acetonitrile with 10mM Ammonium Formate, pH 10.8. The MS analysis alternated between MS and data-dependent MSn scans using dynamic exclusion. The scan range varied slighted between methods but covered 70-1000 m/z.

Raw data was extracted, peak-identified and QC processed using Metabolon's hardware and software. Compounds were identified by comparison to library entries of purified standards or recurrent unknown entities. Metabolon maintains a library based on authenticated standards that contains the retention time/index (RI), mass to charge ratio (m/z), and chromatographic data (including MS/MS spectral data) on all molecules present in the library. Furthermore, biochemical identifications are based on three criteria: retention index within a narrow RI window of the proposed identification, accurate mass match to the library +/- 10 ppm, and the MS/MS forward and reverse scores between the experimental data and authentic standards. The MS/MS scores are based on a comparison of the ions present in the experimental spectrum to the ions present in the library spectrum. The resulting data were further analyzed using the limma-voom Bioconductor package in R to identify differential metabolites with an adjusted pvalue ≤ 0.05[3,4].

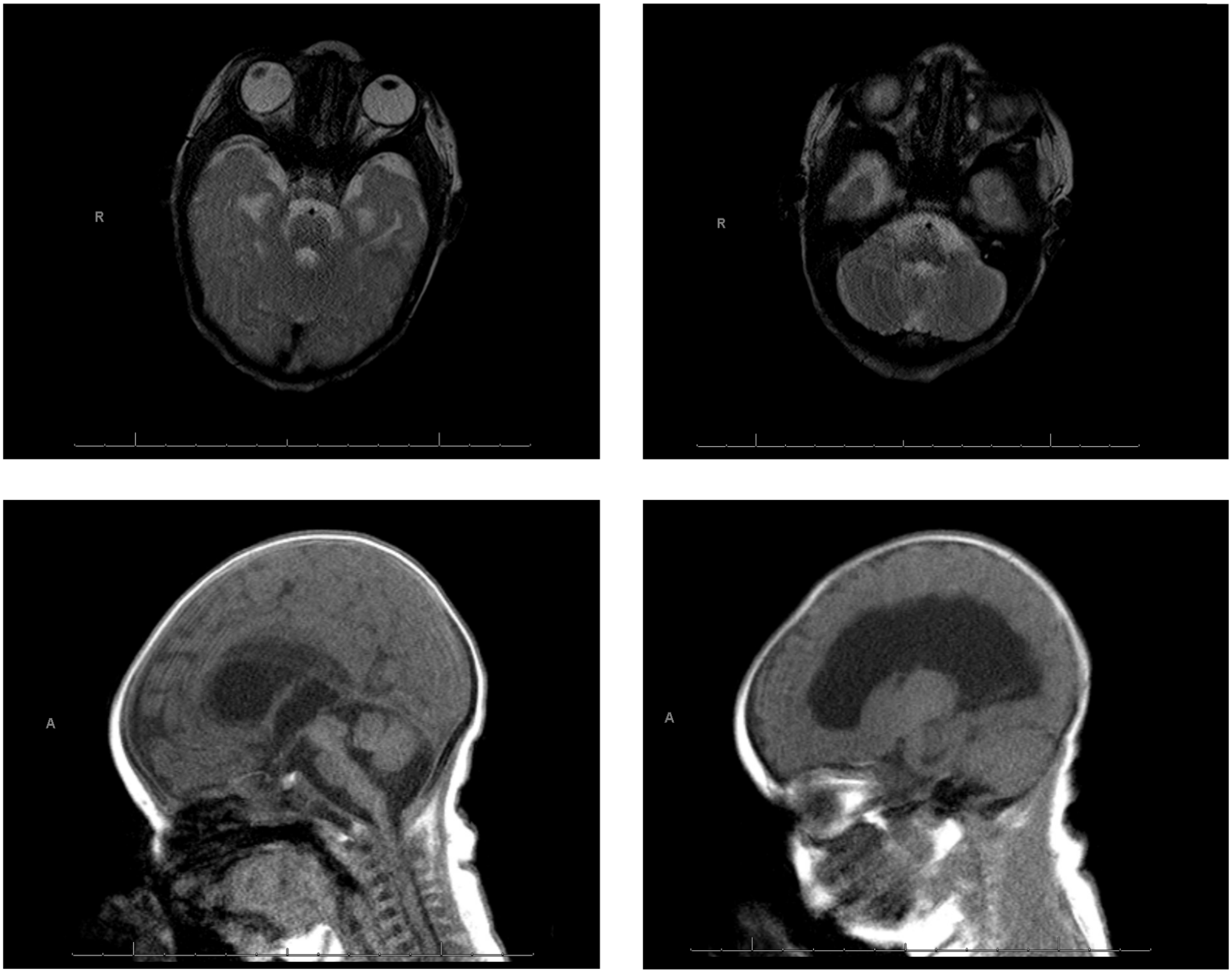


**Supplementary Figure 1. Magnetic Resonance Imaging showed severe white matter loss.** Brain MRI imaging revealed moderate to severe supratentorial ventriculomegaly with irregular margins of the lateral ventricles. The corpus callosum is markedly thinned but does appear to be present. There is a large defect in the septum pellucidum and optic nerves were not well seen.

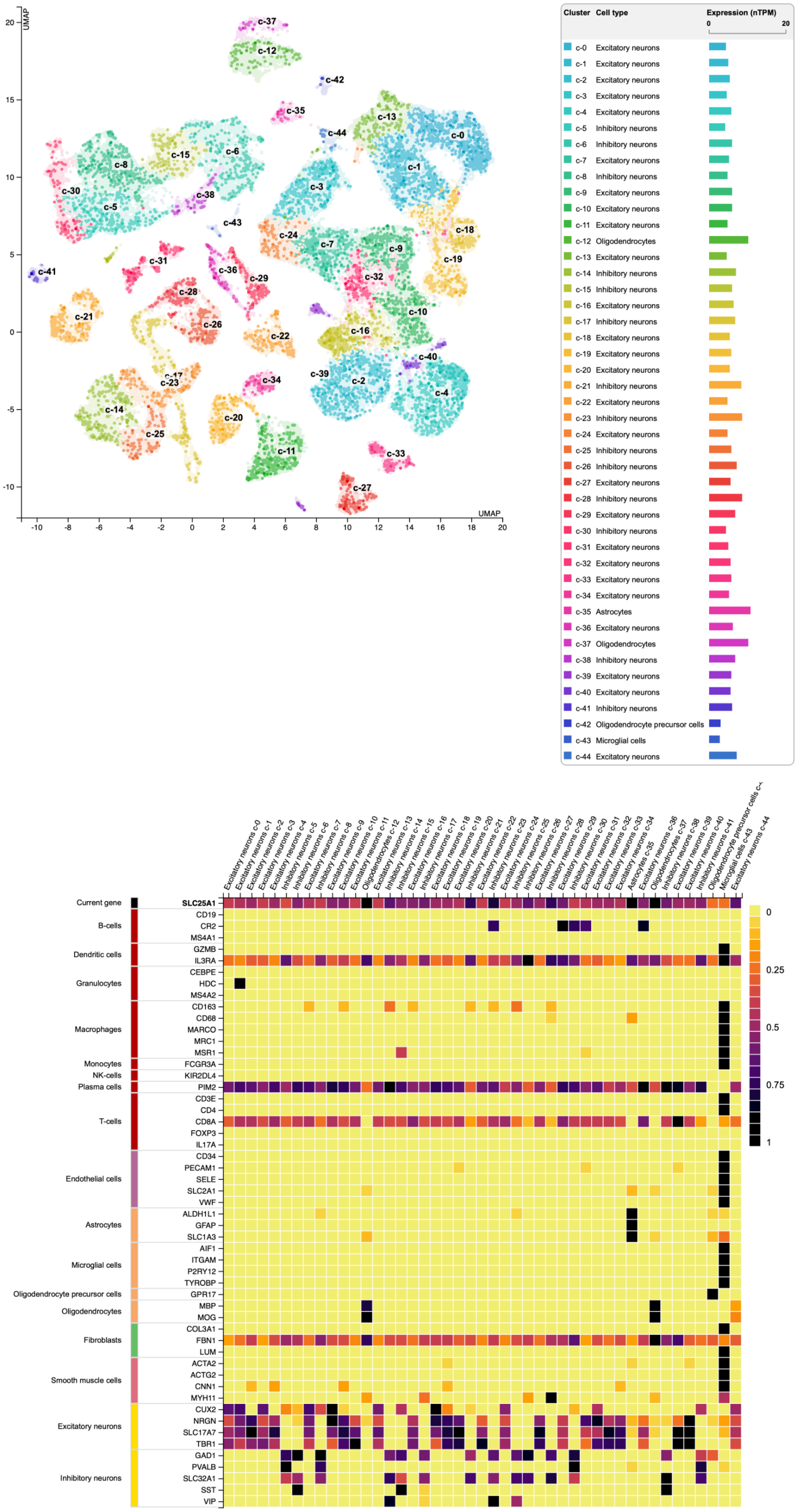


**Supplementary Figure 2. *SLC25A1* is ubiquitously expressed in all neuronal cells.** UMAP analysis derived from The Human Protein Atlas revealed that *SLC25A1* is widely expressed in all neuronal clusters, with the highest expression localized to oligodendrocytes, inhibitory neurons, excitatory neurons and astrocytes.

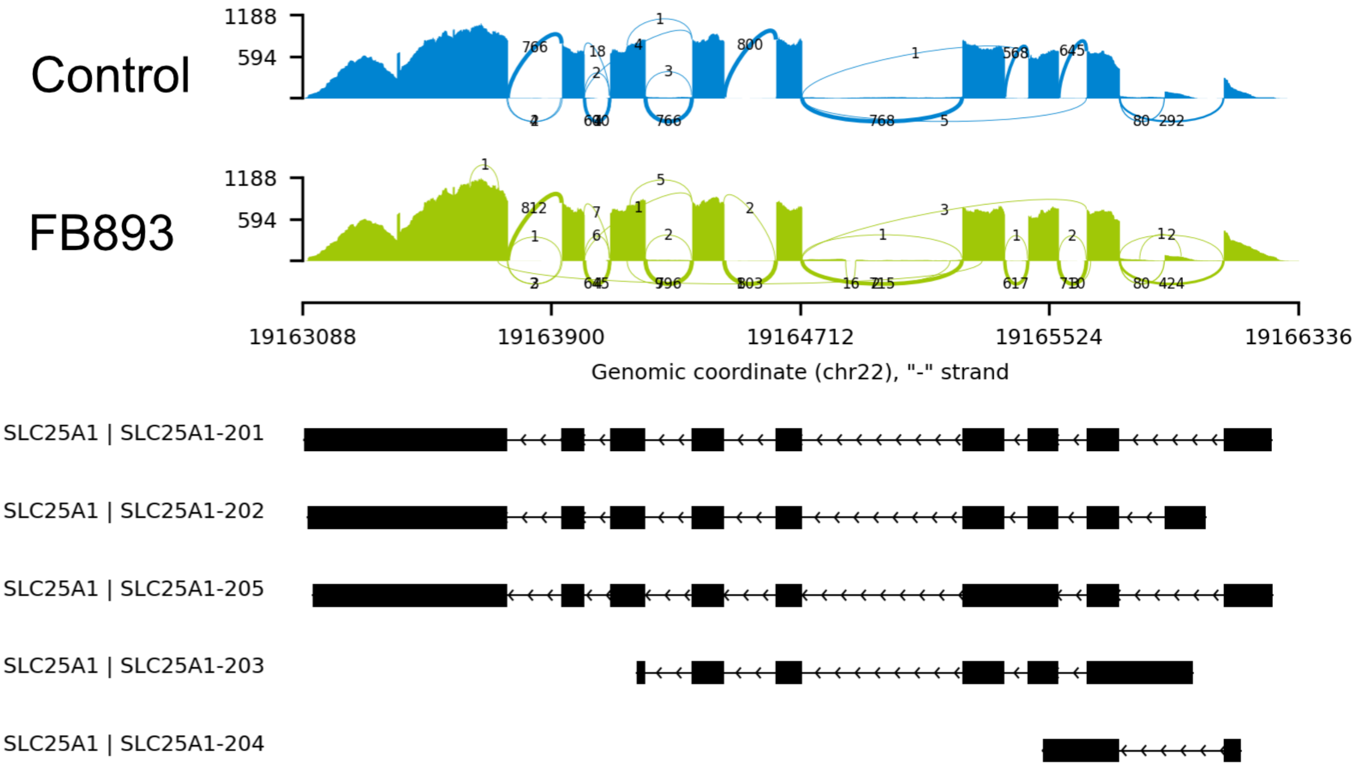


**Supplementary Figure 3. DROP-FRASER and Sashimi plot analysis revealed no evidence of aberrant spliced *SLC25A1* transcripts.** When compared to control fibroblasts, FB893 *SLC25A1* transcripts displayed normal splicing events. The coverage for each alignment track is plotted as a bar graph. Arcs represent splice junctions that connect exons and the number of reads split across the junction (junction depth).

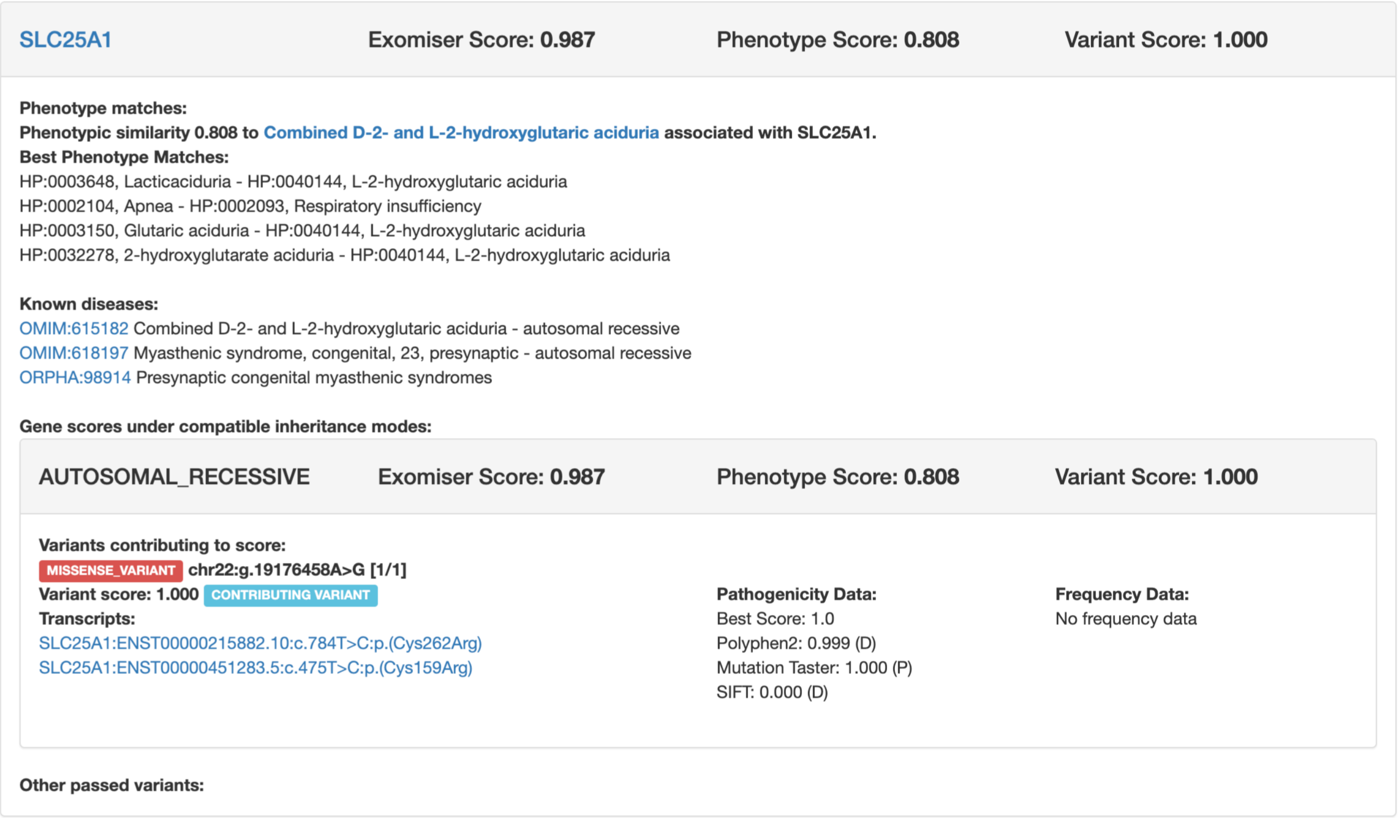

SLC25A1 | Exomiser Score: 0.987 | Phenotype Score: 0.808 | Variant Score: 1.000

**Phenotype matches:**
**Phenotypic similarity 0.808 to Combined D-2- and L-2-hydroxyglutaric aciduria associated with SLC25A1.**
**Best Phenotype Matches:**
HP:0003648, Lacticaciduria - HP:0040144, L-2-hydroxyglutaric aciduria
HP:0002104, Apnea - HP:0002093, Respiratory insufficiency
HP:0003150, Glutaric aciduria - HP:0040144, L-2-hydroxyglutaric aciduria
HP:0032278, 2-hydroxyglutarate aciduria - HP:0040144, L-2-hydroxyglutaric aciduria

**Known diseases:**
OMIM:615182 Combined D-2- and L-2-hydroxyglutaric aciduria - autosomal recessive
OMIM:618197 Myasthenic syndrome, congenital, 23, presynaptic - autosomal recessive
ORPHA:98914 Presynaptic congenital myasthenic syndromes

**Gene scores under compatible inheritance modes:**

AUTOSOMAL_RECESSIVE | Exomiser Score: 0.987 | Phenotype Score: 0.808 | Variant Score: 1.000

**Variants contributing to score:**
MISSENSE_VARIANT **chr22:g.19176458A>G [1/1]**
**Variant score: 1.000** CONTRIBUTING VARIANT
**Transcripts:**
SLC25A1:ENST00000215882.10:c.784T>C:p.(Cys262Arg)
SLC25A1:ENST00000451283.5:c.475T>C:p.(Cys159Arg)

**Pathogenicity Data:**
Best Score: 1.0
Polyphen2: 0.999 (D)
Mutation Taster: 1.000 (P)
SIFT: 0.000 (D)

**Frequency Data:**
No frequency data

**Other passed variants:**

**Supplementary Figure 4. Exomiser analysis of FB893 RNA-seq prioritized *SLC25A1* as the top disease gene candidate.** Analysis of FB893 RNA-seq VCF file revealed a prioritization for the homozygous c.784T>C (p.Cys262Arg) variant in *SLC25A1* (Exomiser Score: 0.987) that is associated with Combined D-2 and L-2-hydroxyglutaric aciduria.

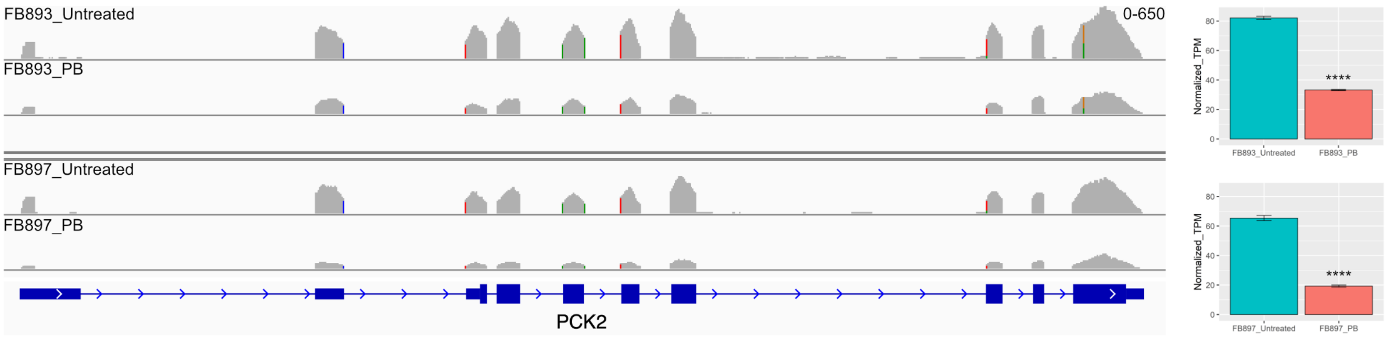


**Supplementary Figure 5. RNA-seq showed a reduction of *PCK2* in phenylbutyrate-treated fibroblasts.** When compared to its untreated state, both phenylbutyrate-treated patient fibroblasts demonstrated a statistically significant reduction in *PCK2*. **** p value ≤ 0.0001.

### Supplementary References


1. Liao, Y., Smyth, G.K., and Shi, W. (2019). The R package Rsubread is easier, faster, cheaper and better for alignment and quantification of RNA sequencing reads. Nucleic Acids Res *47*, e47. 10.1093/nar/gkz114.
2. Liao, Y., Smyth, G.K., and Shi, W. (2014). featureCounts: an efficient general purpose program for assigning sequence reads to genomic features. Bioinformatics *30*, 923-930. 10.1093/bioinformatics/btt656.
3. Law, C.W., Chen, Y., Shi, W., and Smyth, G.K. (2014). voom: Precision weights unlock linear model analysis tools for RNA-seq read counts. Genome Biol *15*, R29. 10.1186/gb-2014-15-2-r29.
4. Ritchie, M.E., Phipson, B., Wu, D., Hu, Y., Law, C.W., Shi, W., and Smyth, G.K. (2015). limma powers differential expression analyses for RNA-sequencing and microarray studies. Nucleic Acids Res *43*, e47. 10.1093/nar/gkv007.
5. Liao, Y., Wang, J., Jaehnig, E.J., Shi, Z., and Zhang, B. (2019). WebGestalt 2019: gene set analysis toolkit with revamped UIs and APIs. Nucleic Acids Res *47*, W199-W205. 10.1093/nar/gkz401.
6. Zhang, B., Kirov, S., and Snoddy, J. (2005). WebGestalt: an integrated system for exploring gene sets in various biological contexts. Nucleic Acids Res *33*, W741-748. 10.1093/nar/gki475.
7. McKenna, A., Hanna, M., Banks, E., Sivachenko, A., Cibulskis, K., Kernytsky, A., Garimella, K., Altshuler, D., Gabriel, S., Daly, M., and DePristo, M.A. (2010). The Genome Analysis Toolkit: a MapReduce framework for analyzing next-generation DNA sequencing data. Genome Res *20*, 1297-1303. 10.1101/gr.107524.110.
8. Smedley, D., Jacobsen, J.O., Jager, M., Kohler, S., Holtgrewe, M., Schubach, M., Siragusa, E., Zemojtel, T., Buske, O.J., Washington, N.L., et al. (2015). Next-generation diagnostics and disease-gene discovery with the Exomiser. Nat Protoc *10*, 2004-2015. 10.1038/nprot.2015.124.
9. Mertes, C., Scheller, I.F., Yepez, V.A., Celik, M.H., Liang, Y., Kremer, L.S., Gusic, M., Prokisch, H., and Gagneur, J. (2021). Detection of aberrant splicing events in RNA-seq data using FRASER. Nat Commun *12*, 529. 10.1038/s41467-020-20573-7.
10. Yepez, V.A., Mertes, C., Muller, M.F., Klaproth-Andrade, D., Wachutka, L., Fresard, L., Gusic, M., Scheller, I.F., Goldberg, P.F., Prokisch, H., and Gagneur, J. (2021). Detection of aberrant gene expression events in RNA sequencing data. Nat Protoc *16*, 1276-1296. 10.1038/s41596-020-00462-5.
11. Frankish, A., Diekhans, M., Ferreira, A.M., Johnson, R., Jungreis, I., Loveland, J., Mudge, J.M., Sisu, C., Wright, J., Armstrong, J., et al. (2019). GENCODE reference annotation for the human and mouse genomes. Nucleic Acids Res *47*, D766-D773. 10.1093/nar/gky955.
12. Dobin, A., Davis, C.A., Schlesinger, F., Drenkow, J., Zaleski, C., Jha, S., Batut, P., Chaisson, M., and Gingeras, T.R. (2013). STAR: ultrafast universal RNA-seq aligner. Bioinformatics *29*, 15-21. 10.1093/bioinformatics/bts635.
13. Yepez, V.A., Gusic, M., Kopajtich, R., Mertes, C., Smith, N.H., Alston, C.L., Ban, R., Beblo, S., Berutti, R., Blessing, H., et al. (2022). Clinical implementation of RNA sequencing for Mendelian disease diagnostics. Genome Med *14*, 38. 10.1186/s13073-022-01019-9.
14. Wagner, A., Wang, C., Fessler, J., DeTomaso, D., Avila-Pacheco, J., Kaminski, J., Zaghouani, S., Christian, E., Thakore, P., Schellhaass, B., et al. (2021). Metabolic modeling of single Th17 cells reveals regulators of autoimmunity. Cell *184*, 4168-4185 e4121. 10.1016/j.cell.2021.05.045.
15. Dobrowolski, S.F., Alodaib, A., Karunanidhi, A., Basu, S., Holecko, M., Lichter-Konecki, U., Pappan, K.L., and Vockley, J. (2020). Clinical, biochemical, mitochondrial, and metabolomic aspects of methylmalonate semialdehyde dehydrogenase deficiency: Report of a fifth case. Mol Genet Metab *129*, 272-277. 10.1016/j.ymgme.2020.01.005.

16. Dobrowolski, S.F., Phua, Y.L., Sudano, C., Spridik, K., Zinn, P.O., Wang, Y., Bharathi, S., Vockley, J., and Goetzman, E. (2022). Comparative metabolomics in the Pah(enu2) classical PKU mouse identifies cerebral energy pathway disruption and oxidative stress. Mol Genet Metab *136*, 38-45. 10.1016/j.ymgme.2022.03.004.
